# Maspin/SerpinB5 is a cytoskeleton-binding protein that regulates epithelial cell shape

**DOI:** 10.1101/2024.09.09.612024

**Authors:** Luiz E. da Silva, Lia M.G. Paim, Ana P. J. Menezes, Julia P. C. da Cunha, Susanne Bechstedt, Nathalie Cella

**Author notes:** Correspondence to Nathalie Cella: ncella.usp.br Av. Prof. Lineu Prestes, 1524, São Paulo, SP, 05508-000, Brazil.

## Abstract

Maspin/SerpinB5 is an abundant and pleiotropic protein mostly expressed by epithelia. Initially described as a tumor suppressor, it has been reported as a regulator of cell adhesion, migration, and invasion. How intracellular Maspin orchestrates these processes is poorly understood. In this study, we utilized Affinity purification-Mass spectrometry (AP/MS) alongside *in vitro* reconstitution assays to establish that Maspin directly interacts with microtubules and microfilaments. Additionally, CRISPR/Cas9-mediated GFP tagging of endogenous Maspin, combined with immunostaining, revealed its localization at the cortical cytoskeleton and the mitotic spindle. Depletion of Maspin by RNAi and CRISPR/Cas9 in three distinct epithelial cell lines disrupts cell-cell adhesion, reorganizes the cytoskeleton and results in upregulation of mesenchymal markers during interphase. In mitotic cells, loss of Maspin induces abnormal cell rounding and rearrangement of cortical F-actin. Moreover, Maspin suppresses microtubule growth *in vitro* and in cells. Collectively, these results demonstrate that Maspin acts at the interface between the cytoskeleton and adhesion sites, directly modulating cell shape and preventing epithelial-mesenchymal transition.

**Summary:** Da Silva et al. report that the non-inhibitory serpin Maspin (SerpinB5), which has long been implicated in the regulation of cell adhesion, migration, invasion and metastasis, directly binds to microfilaments and microtubules *in vitro* and in cells, acting at the interface between the cytoskeleton and adhesion sites.

## Introduction

Maspin (mammary serpin) was initially identified as a tumor suppressor gene in human mammary epithelial cells (Zou et al., 1994). It belongs to the serpin (serine protease inhibitor) family due to structural homology (Law et al., 2006), but compiled data showed it does not inhibit proteases (Pemberton et al., 1995). Maspin is abundant in epithelial cells, it is found in different subcellular compartments and it is extracellularly secreted (Pemberton et al., 1997; Dean et al., 2017). It has pleiotropic biological activities including inhibition of cell proliferation (Machowska et al., 2014), migration (Sheng et al., 1996; Bass et al., 2002; Ravenhill et al., 2010; Qin and Zhang, 2010), invasion (Seftor et al., 1998; Sakabe et al., 2021) and metastasis (Goulet et al., 2011), increase of apoptosis susceptibility (Latha et al., 2005; Mahajan et al., 2019), control of gene transcription (Bailey et al., 2005; Li et al., 2006) and oxidative stress response (Yin et al., 2005). Most studies done in tumor cell lines or animal models report that Maspin has a tumor suppressor activity (Zou et al., 1994; Abraham et al., 2003a; Beltran et al., 2007; Shao et al., 2008; Dzinic et al., 2017a). However, clinical data have challenged this concept, as Maspin was found to be expressed in advanced thyroid (Ito et al., 2004), gastric (Kim et al., 2005), colorectal (Snoeren et al., 2013) and breast cancers (Ahn et al., 2002). This divergence may be explained in part by differences in the experimental approach and model system. In any case, it is becoming well accepted that Maspin has a dual nature, sometimes restraining cancer progression, other times promoting it. We and others have shown that addition of recombinant Maspin to the cell medium results in modulation of cell adhesion and motility (Sheng et al., 1996; Seftor et al., 1998; Cella et al., 2006). This process depends on the interaction between Maspin, the uPA/uPAR complex and β1-integrin (Yin et al., 2006; Endsley et al., 2011). In support of these findings, Maspin has been detected in the extracellular space *in vivo*, in a tumor context (Tian et al., 2020). Despite these observations, it is well known that Maspin is predominantly an intracellular protein both in normal and tumor cells that retain its expression (Thul et al., 2017). The role of intracellular Maspin has been investigated mostly by Maspin ectopic gene expression in tumor cells that lost its expression(Zou et al., 1994; Jiang et al., 2002; Abraham et al., 2003b; Latha et al., 2005; Zhang et al., 2005). In cells, Maspin re-expression inhibited cell migration and invasion (Zou et al., 1994; Abraham et al., 2003a; Ravenhill et al., 2010), similar to observations done in tumor cells which have been exposed to extracellular Maspin (Sheng et al., 1996; Bass et al., 2002; Yin et al., 2006; Bass et al., 2009; Ravenhill et al., 2010; Endsley et al., 2011). *In vivo*, however, the observations were divergent, similar to the clinical findings in this field - when breast and lung tumor cells ectopically expressing Maspin were either injected or transplanted into mice, tumors were smaller and less aggressive (Shi et al., 2001; Nam and Park, 2010). Opposingly, a similar strategy used in pancreatic tumor cells favored tumor progression (Tian et al., 2020). Our experimental approach is different from others in two important aspects - we used non-transformed cell lines (Tamazato Longhi and Cella, 2012; Longhi et al., 2021) and the normal mouse mammary gland development (Longhi et al., 2016) as model systems to investigate how Maspin mechanistically regulates cell function. Our objective was to investigate the relation between intracellular Maspin and cytoskeleton proteins. Here, we uncover that Maspin modulates cytoskeleton organization by directly interacting with actin filaments and microtubules. Maspin deletion leads to reorganization of cortical actin, microtubules and disruption of cell-cell adhesion, triggering epithelial-mesenchymal transition. Since cytoskeleton networks drive cell adhesion, shape and integrate biochemical signals between the cytoplasm and the nucleus, these data hold promise for resolving and reconciling divergent observations in health and disease.

## Results and Discussion

### AP/MS analysis reveals that endogenous Maspin interacts with the cytoskeleton

We initially selected non-transformed epithelial cell lines in order to study the function of the underexplored endogenous Maspin. Among them, immortalized human and mouse mammary epithelial cells (MCF-10A and EpH4, respectively) and immortalized human keratinocyte cells (HaCaT) express endogenous Maspin, as observed by Western blot using an anti-Maspin monoclonal antibody (Fig. 1A). We have previously used IP/MS to capture endogenous Maspin interacting proteins from MCF-10A cell lysates. In our analysis, Maspin consistently interacts with α-tubulin (Longhi et al., 2021). Given the essential role of microtubules throughout the cell cycle, we explored Maspin’s possible interacting partners during interphase and cell division. To this end, we arrested MCF-10A cells at G2/M transition using nocodazole (Fig. 1B), which could be confirmed by rounded morphology compared to DMSO-treated cells (Fig. 1C). Cell lysates were incubated with recombinant GST and GST-Maspin baits and further purified. The eluates were separated by SDS-PAGE and visualized by silver staining. GST- Maspin and GST were promptly detected (Fig. 1D, arrows) as well as different co-eluted material. Proteomics analysis identified interacting proteins exclusively found or enriched in GST-Maspin baits as represented by heatmap according to LFQ Fold Change values (Fig. 1E). String database and Gene Ontology integration indicated that most of the captured interacting proteins are cytoskeleton components (Fig. 1F). Among them, we identified Plectin (PLEC), a plakin family member that links the three main components of the cytoskeleton to adhesion sites (Jefferson et al., 2004); α-actinin-4 (ACTN4), an actin cross-linking protein (Sjöblom et al., 2008), and subunits of the non-muscle myosin II (MYH9 and MYL6), which controls actomyosin contractility (Rosenblatt et al., 2004). For both conditions (DMSO vs. Nocodazole), we consistently detected two interacting proteins - Tubulin Alpha 1c (TBA1C) and Actin Gamma (ACTG) - suggesting that Maspin potentially interacts with cytoskeleton subunits as well. We further confirmed Maspin-actin and Maspin-tubulin interactions in non-arrested condition (no Nocodazole) using pull-down followed by Western blot against actin and tubulin (Fig. 1G). Interestingly, recombinant Maspin captured endogenous Maspin, indicative of possible self-association, as previously observed (Liu et al., 1999; Al-Ayyoubi et al., 2004).

**Figure 1.**
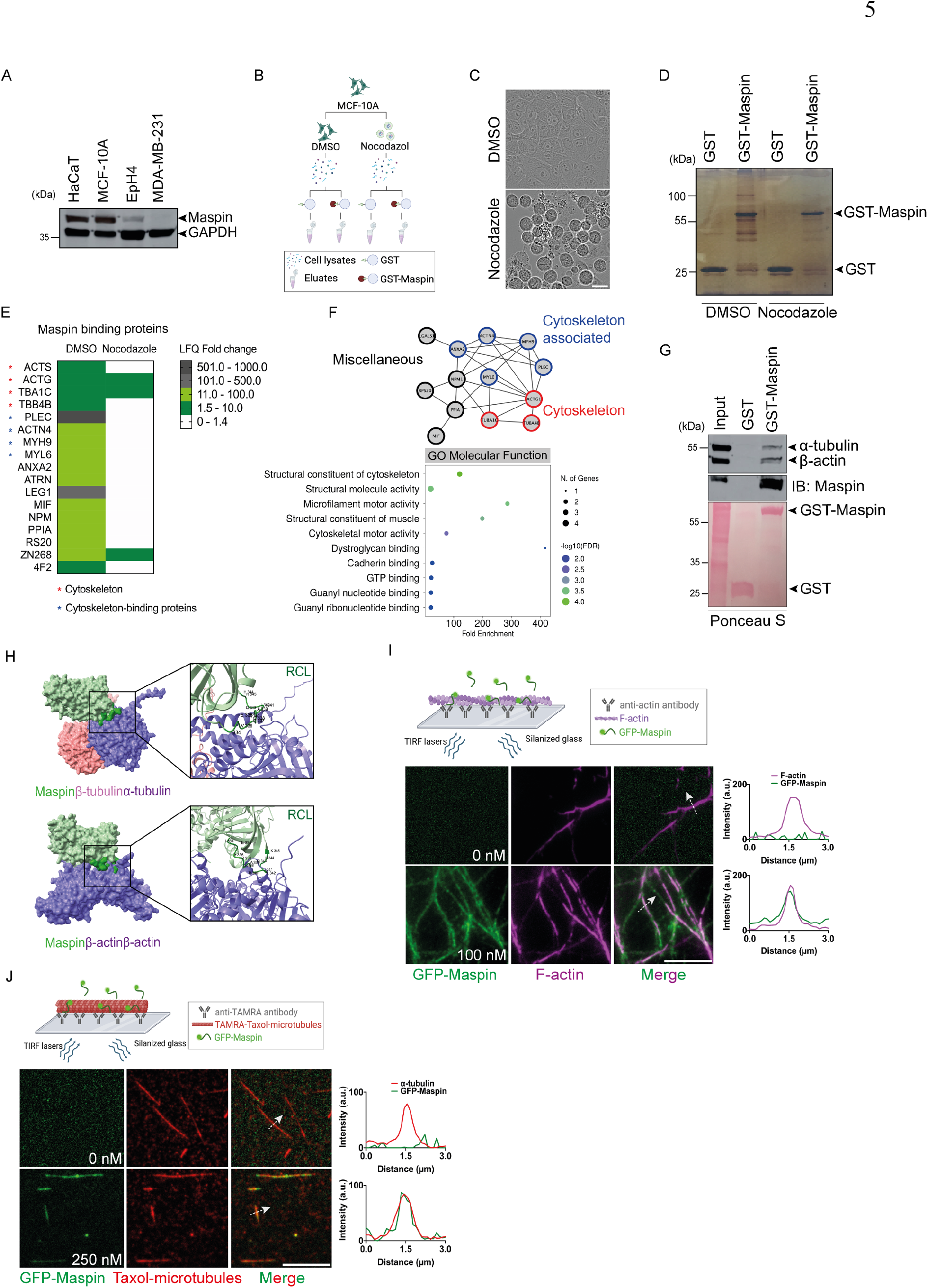
Maspin directly interacts with the cytoskeleton. (A) Maspin protein expression in HaCaT, MCF-10A, EpH4 and MDA-MB-231 cells analyzed by Western blot with an anti-Maspin antibody; **(B)** Workflow of final pull-down eluates submitted to proteomic analysis. MCF-10A cell lysates previously treated with Nocodazole or DMSO were incubated with recombinant GST-Maspin or GST-only negative control as baits; **(C)** Bright-field image of Nocodazole-arrested (rounded morphology) or DMSO-treated (flatten morphology) MCF-10A cells. Scale bar: 25 µm; **(D)** GST- and GST-Maspin pull-down eluates were analyzed by SDS-PAGE followed by silver staining. Eluates were submitted to proteomic analysis; **(E)** Heatmap for interacting proteins exclusively identified or enriched in GST- Maspin samples over GST-only negative control according to normalized Label Free Quantification (LFQ) Fold Change values. Maspin is not depicted (see Suppl. Table 1); **(F)** Protein-protein interaction network generated by String for all hits found in (E). Black edges represent reported physical interactions and nodes were colored according to protein class. Top 10 Molecular Function terms ranked by FDR score in Gene Ontology analysis (lower); **(G)** GST-pull-down (same as in D), followed by Western blot against α-tubulin, β-actin and Maspin (upper membranes). GST-only and GST-Maspin baits were visualized by Ponceau staining prior to Western blot (lower membrane); **(H)** Maspin complexed to αβ-tubulin heterodimer (upper) or with β-actin homodimer (lower) according to AlphaFold multimer. Maspin RCL (Reactive Center Loop) region is highlighted in dark green; **(I)** Schematic representation of F-actin *in vitro* reconstitution assay (upper). Pre-assembled AlexaFluor-labeled F-actin in the presence of purified GFP-Maspin (100 nM) was visualized by TIRF microscopy (lower); (**J**) Schematic representation of Taxol-stabilized microtubules *in vitro* reconstitution assay (upper). Pre-assembled TAMRA-labeled microtubules in the presence of purified GFP-Maspin (250 nM) were visualized by TIRF microscopy (lower). Scale bar: 10 µm.

As we captured other cytoskeleton-binding proteins, we reasoned that Maspin could be itself a cytoskeleton interacting protein. We then checked for a presumed direct binding of Maspin to actin and tubulin using AlphaFold multimer (Fig. 1H). Modeling predictions suggested that Maspin interacts through its Reactive Center Loop motif (RCL, aa 334-345) with the surface of α-tubulin (aa 401-450) on the αβ-tubulin heterodimer. The RCL region was also predicted to interact with a region on the β- actin homodimer (aa 348-355). Next, we validated the direct interaction between Maspin and microfilaments and microtubules using total internal reflection fluorescence (TIRF) microscopy-based *in vitro* reconstitution assays. To this end, we generated an N-terminal fluorescently-tagged recombinant Maspin and purified it using a His-StrepII double-tag purification strategy (Suppl. Fig. 1A) (McAlear and Bechstedt, 2022). Tagged Maspin was added to pre-assembled F-actin and Taxol-stabilized microtubules and analyzed by TIRF microscopy. Purified recombinant Maspin promptly bound to F- actin (Fig. 1I) and to stabilized microtubules (Fig. 1J) in the nanomolar range *in vitro*. Collectively, these data indicate that Maspin is a cytoskeleton-binding protein.

### Maspin localizes to the cytoskeleton of epithelial cells

Previous studies reported that Maspin co-localizes with β1-integrin and distributes along cortical F-actin (Cella et al., 2006). Moreover, it was captured from the cell cortex of interphase and mitotic HeLa cells in a large-scale proteomic study (Vadnjal et al., 2022). In monolayer cell culture, MCF-10A cells are known to spontaneously shift their arrangement (Debnath et al., 2003) and nature (Xu et al., 2017) according to confluency. At low to medium confluency, they pack into clusters, exhibiting discrete lamellipodial protrusions, whereas typical cobblestone epithelial organization is adopted at high confluency (Debnath et al., 2003). In order to track Maspin spatiotemporal distribution in live MCF-10A cells, we fluorescently tagged the C-terminal and N-terminal regions of endogenous Maspin by a CRISPR/Cas9-mediated knock-in strategy (Suppl. Fig. 1B). Western blot analysis for all knock-in stable pools with anti-GFP and anti-Maspin antibodies indicated the expected molecular weight bands for fusion proteins (Suppl. Fig. 1C). As repeatedly observed by immunostaining (Bodenstine et al., 2014; Longhi et al., 2016), endogenously tagged Maspin is highly abundant and ubiquitously distributed in a diffuse pattern, but locates mostly to the cytoplasm of all three knock-in cells, as observed by fluorescence live cell imaging (Suppl. Fig. 1D). Given that we did not observe substantial changes in Maspin subcellular localization among all three knock-in pools and due to a higher editing efficiency observed for the N1-terminus-tagged Maspin (Suppl. Fig. 1C), we selected this cell population to perform our downstream assays (hereafter named GFP-N1-Maspin knock-in).

To evaluate GFP-N1-Maspin subcellular localization in cells, we monitored the fluorescence within 24 h post-seeding by live imaging. Notably, Maspin localizes at the leading edge of spreading cells and to lamellipodia of collectively migrating cells (Fig. 2A, left panel, and video 1). As confluency increases, it becomes mostly confined to cytoplasm, adopting a diffuse pattern (Fig. 2A, middle panel). Unexpectedly, Maspin also accumulates at the spindle zone of cells undergoing mitosis (Fig. 2A, right panel), which has not been previously reported. Because Maspin is highly abundant and epifluorescence usually hampers an accurate visualization of its subcellular localization, we performed confocal immunofluorescence analysis of MCF-10A WT cells with an anti-Maspin monoclonal antibody. Sections along the z-axis indicated a similar distribution pattern - Maspin is found in the cytoplasm and it is also spatially distributed along microfilaments and microtubules, either colocalizing at the cytoskeleton of lamellipodial projections of epithelial cells or co-distributing at the cell cortex (Fig. 2B, upper and middle panels). In addition, it accumulates at the spindle zone and sparsely decorates the microtubules of the mitotic spindle (Fig. 2B, lower panel), which was confirmed in HaCaT cells (Suppl. Fig. 1E). Given that Maspin subcellular localization is susceptible to cell-cell contact regulation (Longhi et al., 2021), we suspect it may partition dynamically between microfilaments and microtubules according to mechanical and biochemical signals emanating from cell-cell adhesion sites. A similar mechanism has been described for the Hippo-yes-associated protein (Yap) (Das et al., 2016; Nardone et al., 2017) or β-catenin (Dietrich et al., 2002; Sen et al., 2022), which are controlled by cell adhesion and cytoskeleton network and also exhibit nucleocytoplasmic shuttling. In addition, Maspin detection at the spindle zone of mitotic cells further reinforces the concept that it dynamically regulates both microfilaments and microtubules in epithelial cells.

**Figure 2.**
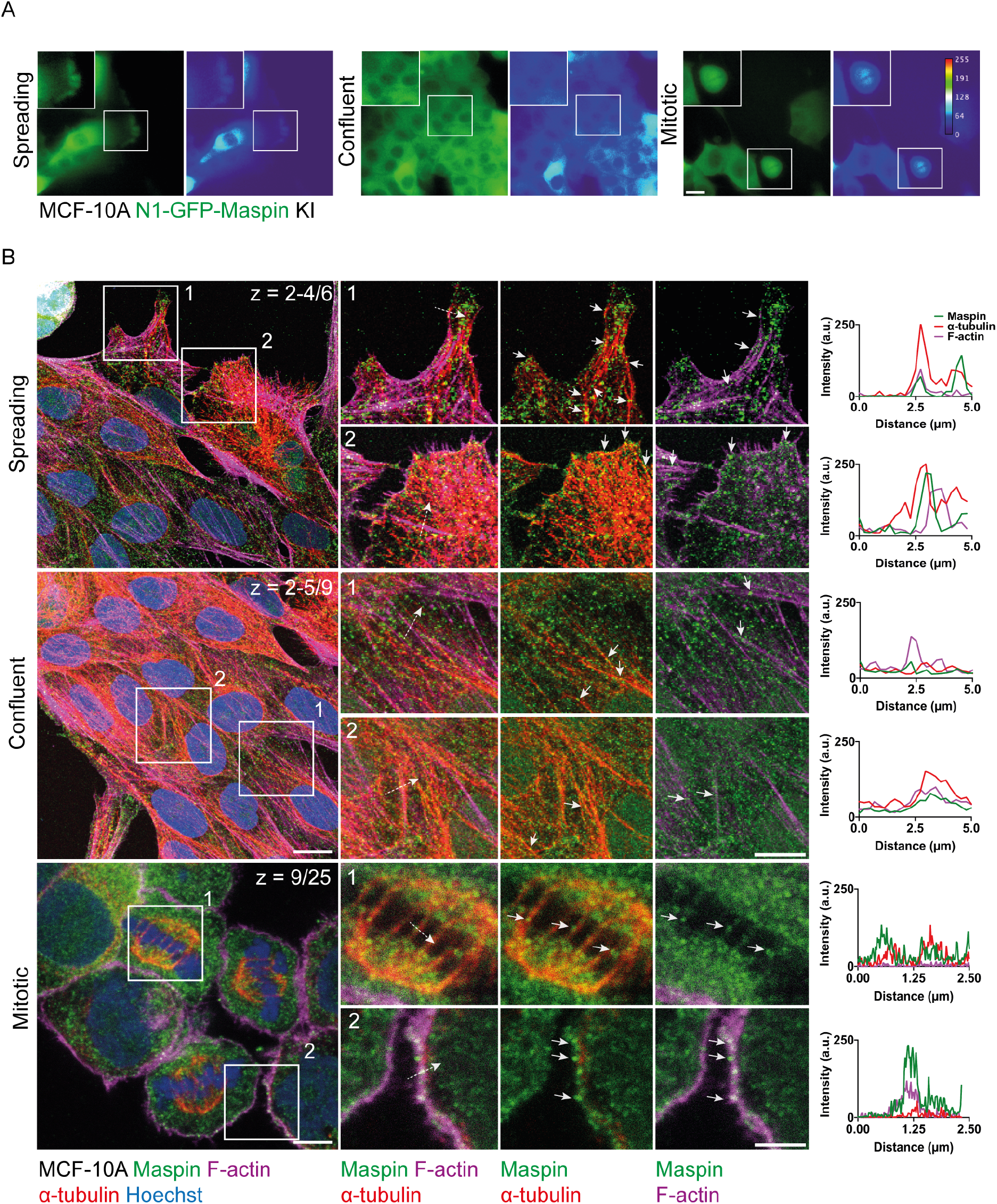
Endogenous Maspin distributes along the cytoskeleton. (A) MCF-10A cells expressing endogenously tagged Maspin observed by fluorescence live imaging. Maspin is promptly detected at the leading edge of spreading cells, it becomes ubiquitous at the cytoplasm of confluent cells and accumulates at the spindle zone of mitotic cells. Scale bar: 10 µm; **(B)** Confocal immunofluorescence analysis in fixed MCF-10A cells. Co-staining of Maspin, α-tubulin and F-actin confirmed its localization along microfilaments and microtubules of the cell cortex and mitotic spindle. Filled white arrows indicate Maspin along microfilaments and microtubules. Dashed arrows refer to fluorescence intensity plots graphics on the right side. Scale bar: 10 µm.

### Loss of endogenous Maspin induces spatial reorganization of the cytoskeleton

The contribution of Maspin to *in vivo* lethality has been addressed by different studies. Although divergent data brought controversy to the field (Teoh et al., 2014), it became established that Maspin is essential to embryogenesis and tumor suppression (Gao et al., 2004; Dzinic et al., 2017b; Whisstock and Bird, 2017; Zhang et al., 2017b). Depletion of endogenous Maspin in epithelial cells has been typically carried out using RNAi (Dean et al., 2017; Sakabe et al., 2021) and, to our knowledge, Maspin knock- out clones have not been previously reported. To better understand the contribution of Maspin to cytoskeleton organization, we established Maspin knock-out (KO) clones using CRISPR/Cas9-genome editing in MCF-10A and HaCaT cells (Suppl. Fig. 1F). Sequencing analysis of edited regions confirmed the presence of several insertions and deletions (Suppl. Fig. 1G), which were quantitatively described by Inference of CRISPR Edits (ICE) algorithm (Suppl. Fig. 1H) (Conant et al., 2022). Subsequently, Western blot analysis verified the complete absence of the protein in all KO clones. (Suppl. Fig. 1I). We thus selected the MCF-10A KO clones C10 and D9 and the HaCaT KO clone C5 to further investigate Maspin function (Suppl. Fig 1H).

We first co-stained F-actin and microtubules to gauge for cytoskeletal changes in our MCF-10A KO cells. Strikingly, confocal z-stack sections indicated that cortical F-actin thickness is reduced, whereas there is a slight increase in stress fibers compared to WT cells (Fig. 3A and B). They also exhibit increased cytoplasmic microtubule density at z-stack basal slices compared to their predominant cortical localization in MCF-10A WT cells (Fig. 3A and C). To rule out the possibility that phenotypic changes associated to Maspin depletion were due to clonal artifacts as previously speculated (Teoh et al., 2014), we also transiently knocked-down Maspin in MCF-10A cells using commercially available siRNA sequences with low (siMaspin #60) and intermediate efficiencies (siMaspin #95) (Suppl. Fig. 2A) and in EpH4 cells stably expressing shMaspin (Suppl. Fig. 2C). Similar cytoskeleton rearrangements were observed in MCF-10A siMaspin cells and EpH4 stably expressing shMaspin (Suppl. Fig. 2B and D, respectively). Of note, transient knock-down of Maspin in MCF-10A cells led to vimentin redistribution from the cell cortex to the cytoplasm, where it becomes more fragmented (Suppl. Fig. 2B).

**Figure 3.**
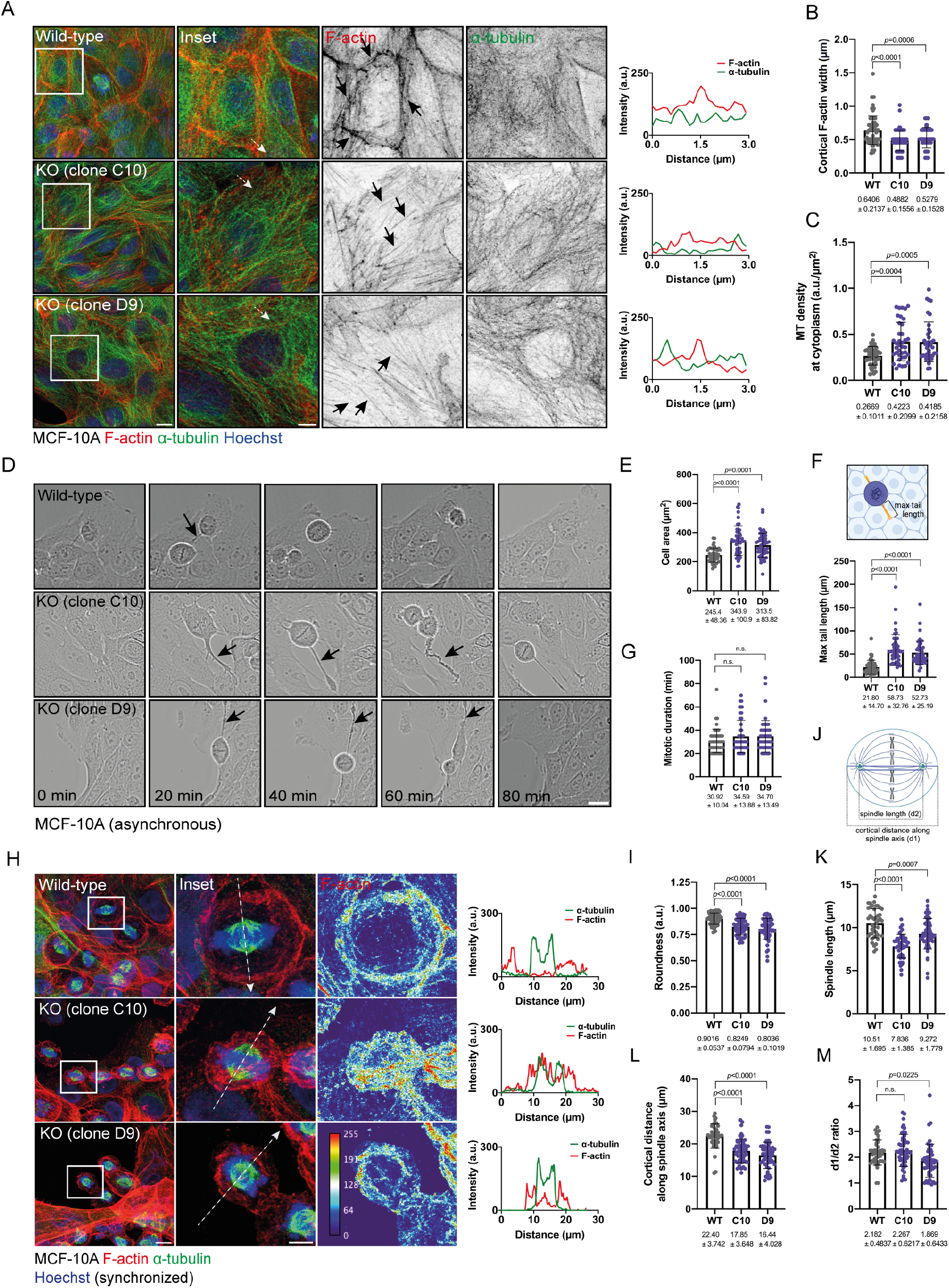
Loss of endogenous Maspin promotes cytoskeleton rearrangements in epithelial cells. **(A)** Fluorescence analysis of F-actin and α-tubulin in MCF-10A KO versus MCF-10A WT cells during interphase. Cortical F-actin (WT) and stress fibers (KO C10 and D9) are indicated by black arrows. Scale bar: 10 µm; **(B)** Cortical F-actin thickness measured for MCF-10A WT cells (n = 74 cells), KO C10 (n = 62 cells) and KO D9 (n = 64 cells); **(C)** Cytoplasmic microtubule density at basal slices of MCF-10A WT (n = 43 cells), KO C10 (n = 38 cells) and KO D9 (n = 39 cells); **(D)** Live imaging of MCF-10A WT *versus* KO cells upon mitotic entry. Black arrows indicate retraction tails. Scale bar: 25 µm; **(E)** Cell area prior to mitotic rounding WT (n = 49 cells), KO C10 (n = 49 cells), KO D9 (n = 49 cells); **(F)** Maximal tail length for WT (n = 50 cells), KO C10 (n = 48 cells) and KO D9 (n = 50 cells) cells during mitotic progression; **(G)** Mitotic duration for WT (n = 50 cells), KO C10 (n = 48 cells) and KO D9 (n = 50 cells). Cells were imaged at interval frames of 5 min; **(H)** MCF-10A cells released after nocodazole blockage and co-stained for F-actin, α-tubulin and nuclei. Heatmaps indicate distribution of F-actin fluorescence intensity in both MCF-10A WT and KO cells. Scale bar: 10 µm and 5 µm (inset); (**I**) Cell roundness was measured for MCF-10A WT (n = 59 cells), KO C10 (n = 53 cells) and KO D9 (n = 53 cells) at metaphase; (**J**) Representation of mitotic measurements (d1 and d2). (**K**) Spindle length (d2) was measured for WT (n = 43 cells), C10 (n = 48 cells) and D9 (n = 58 cells); (**L**) Cortical distance along spindle axis (d1) measured for WT (n = 43 cells), KO C10 (n = 52 cells), KO D9 (n = 52 cells) cells at metaphase; **(M)** Cortical distance along spindle axis to spindle length ratio (d1/d2) measured for WT (n = 43 cells), C10 (n = 52 cells) and D9 (n = 50 cells) at metaphase. n.s. = not significant.

Since Maspin was detected at the spindle zone (Fig. 2B and Suppl. Fig. 1E) and it interacts with actin and tubulin in nocodazole-arrested cells, we assessed the mitotic population of asynchronous MCF- 10A WT and KO cells. Upon mitotic entry, cells partially loose adhesion to the substrate and acquire a rounded morphology (Jones et al., 2019). Thus, we followed the morphology of rounded cells by bright- field live imaging (Fig. 3D). Interestingly, KO cells exhibit an increased area of attachment (Fig. 3E) and longer retraction tails that remain connected to the neighboring cells upon mitotic entry (Fig. 3D, video 2). These abnormal retraction tails are usually bipolar, but asymmetric in length (Fig. 3F). Overall, we observed a slight mitotic delay, which may be related to the increased adhesion area and longer retraction tails (Fig. 3G). These changes, however, do not result in cell death following mitosis (video 2). Next, we arrested MCF-10A KO cells with nocodazole at mitotic entry and monitored their progression after release from Nocodazole-block. F-actin and microtubule staining indicated substantial morphological changes in mitotic cells at metaphase (Fig. 3H). Notably, metaphasic MCF-10A KO cells display reduced roundness (Fig. 3I), with F-actin often randomly distributed at the cell cortex. We then measured mitotic parameters (Fig. 3J) and observed that Maspin KO cells display decreased spindle length (Fig. 3K), which could also be seen in asynchronous MCF-10A-siMaspin cells (Suppl. Fig. 2F) and in thymidine-synchronized EpH4-shMaspin cells (Suppl. Fig. 2G). Because the spindle distance scales according to cell size (Krüger and Tran, 2020) and our synchronized KO cells exhibit reduced size, we measured the cortical distance along the spindle axis (Fig. 3L, d1) to check if reduction in spindle length (Fig. 3K, d2) was due to the reduced cell size. This is true for KO C10 cells, but not for KO D9 cells (Fig. 3M, d1/d2), suggesting that additional factors may contribute to the reduction in spindle length in Maspin KO cells. In addition, occasional distortion of spindle positioning was also observed in MCF-10A KO D9 cells (Fig. 3H, lower panel). Overall, Maspin depletion promotes reorganization of the cell cortex and spindle abnormalities under mitotic synchronization. As Maspin interacts with both microfilaments and microtubules, our KO model system cannot distinguish whether these phenotypes are a consequence of Maspin acting separately on either one or the other. Of note, we also captured subunits of myosin II (MYL6 and MYH9) in our AP/MS analysis and it has been demonstrated that actomyosin controls spindle positioning (Rosenblatt et al., 2004). As Maspin regulates RhoGTPase activity in different model systems (Odero-Marah et al., 2003; Shi et al., 2007), we suspect that its loss at the cell cortex may modify cell contractility, phenotypically resembling an imbalance in Rho/ROCK/myosin II signaling (Maddox and Burridge, 2003; Chauhana et al., 2011).

### Maspin negatively regulates microtubule growth *in vitro* and in cells

Although some evidence suggested that Maspin regulates the actin cytoskeleton (Al-Mamun et al., 2016), no previous report addressed its role on microtubules. Therefore, we decided to investigate whether Maspin regulates microtubule growth dynamics in cells. For this purpose, we first analyzed how microtubules recover from nocodazole-induced depolymerization in Maspin-depleted cells. Surprisingly, microtubule assembly appeared to be faster in MCF-10A Maspin KO (Fig. 4A and B), in MCF-10A siMaspin cells (Suppl. Fig. 2H) and in EpH4 shMaspin cells (Suppl. Fig 2I) after nocodazole washout. At 3.5min post-Nocodazole washout, microtubules in Maspin KO cells appeared to regrow more radially and display a cytoplasmic organization similar to that observed in non-treated Maspin KO cells (Fig. 3A), whereas microtubule regrowth in WT cells was much less prominent (Fig. 4A and B). Next, we assessed microtubule dynamics by transiently transfecting MCF-10A Maspin KO cells with a construct encoding EB3-StayGold, a microtubule-plus-end-binding tracker (Fig. 4C, video 3). In these cells, we observed an increase in microtubule growth rates (Fig. 4D) and a higher number of comets per area (Fig. 4E) as compared to MCF-10A WT cells. Interestingly, maximal intensity projections over time indicated that EB3 tracks adopt a crossed pattern, as indicated by the lower anisotropy score for both KO cells (Fig. 4F), which is consistent with the radial growth pattern observed in our microtubule regrowth assays. Taken together, these data suggest that Maspin suppresses microtubule growth, and spatially reorganizes the microtubules of epithelial cells.

**Figure 4.**
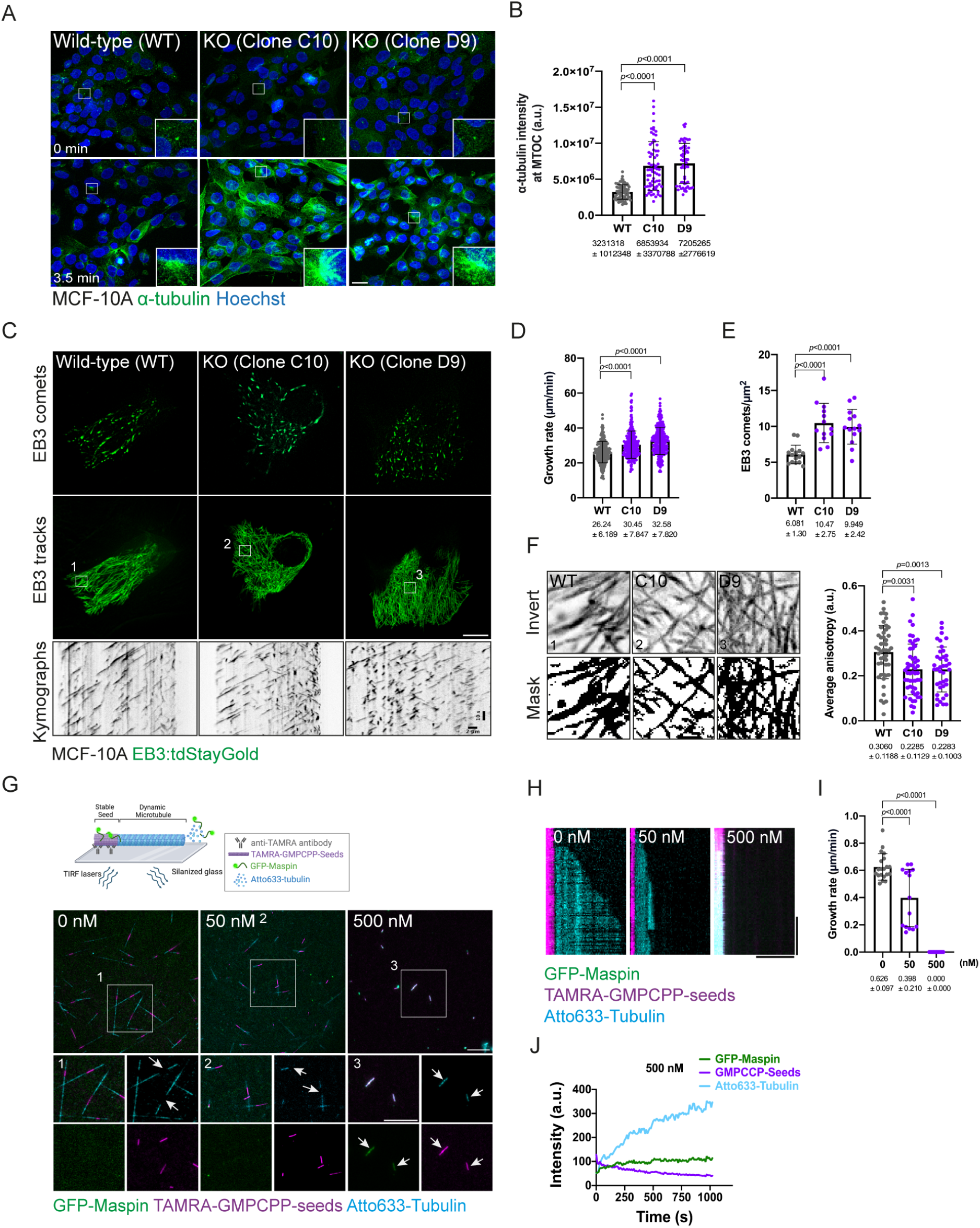
Maspin regulates microtubule dynamics *in vitro* and in cells. (A) Microtubule recovery from nocodazole-induced depolymerization in MCF-10A WT, KO C10 and KO D9 cells. Cells were released and fixed after 0 min and 3.5 min and stained for α-tubulin; **(B)** Quantification of α-tubulin intensity around each microtubule-organizing center (MTOC) after 3.5 min of nocodazole washout in WT (n = 77 cells), KO C10 (n = 70 cells) and KO D9 (n = 59 cells); **(C)** Live imaging of MCF-10A WT, KO C10 and KO D9 cells expressing EB3-StayGold (upper panel). Representative EB3 maximal intensity projections (tracks) (middle panel) and kymographs (lower panel); **(D)** Microtubule growth rate (µm/min) in WT (n = 263 comets, 20 cells), KO C10 (n = 251 comets, 19 cells), KO D9 (n = 261 comets, 17 cells); **(E)** Average of EB3 comets per area in WT (n = 17 cells), KO C10 (n = 13 cells) and KO D9 (n = 17 cells); **(F)** Average anisotropy in WT (n = 14 cells), KO C10 (n = 17 cells) and KO D9 (n = 13 cells); **(G)** Schematic representation of dynamic microtubules using an *in vitro* reconstitution assay (upper). Pre-assembled TAMRA-labeled GMPCPP-seeds in the presence of Atto633-tubulin and purified GFP-Maspin at different concentrations were visualized by TIRF microscopy (lower). Scale bar: 10 µm; **(H)** Representative kymographs extracted from dynamic microtubules in (G). Scale bar: 5 µm (horizontal), time: 5 min (vertical); **(I)** Plot of microtubule growth rates as a function of GFP-Maspin concentration with 8 µM tubulin. Control (0 nM; n = 20 microtubules), GFP-Maspin (50 nM; n = 15 microtubules) and GFP-Maspin (500 nM; n = 15 microtubules); **(J)** Fluorescence intensity over time for GMPCPP-seeds exposed to GFP-Maspin (500 nM). Maspin binds to the seeds and tubulin is increasingly recruited to them.

Next, we wondered if the observed suppression of microtubule growth was an indirect effect of Maspin, or a direct consequence of Maspin interaction with microtubules. As we have shown that Maspin directly interacts with stabilized microtubules (Fig. 1J), this prompted us to probe for the direct effect of Maspin on microtubule growth using a TIRF-based *in vitro* reconstitution assay. To this end, we assembled dynamic microtubules in the presence of stable GMPCPP-seeds and purified recombinant Maspin. Notably, we noticed a significant suppression of microtubule growth when Maspin was added at low concentration (50 nM) (Figure 4G-I, video 4). This suppressive effect increases at high concentration (500 nM) (Figure 4G-I, video 4), where Maspin also readily decorates and recruits free tubulin to GMPCPP-seeds (Figure 4G, white arrows, and J). We could not observe, however, the binding of Maspin to dynamic microtubules. Therefore, we conclude that the effect of growth suppression in cells is a direct effect of Maspin’s impact on microtubule growth. Similar as in cells, we also do not observe a strong interaction of Maspin with the growing microtubule lattice. Growth suppression may therefore result from transient interactions of Maspin with the growing microtubule end. This mechanism is supported by the strong interaction of Maspin with GMPCPP-stabilized microtubules (Figure 4 G and H), which is a proxy for the GTP cap at microtubule ends (Roostalu et al., 2020). Further experiments are necessary to determine how Maspin mechanistically interacts with free tubulin or microtubules to promote their destabilization. Interestingly, in addition to microtubules, several microtubule-associated proteins (MAPs) interact with F-actin (Pimm and Henty-Ridilla, 2021). Conversely, actin-binding proteins are also able to directly interact with microtubules and regulate their dynamics (Villari et al., 2015). In the case of Maspin, because both interactions appear to predominantly occur through the same RCL motif, we speculate that they may take place in a mutually exclusive manner, as reported for some MAPs (Alberico et al., 2016; Wu et al., 2022). These molecules bind to F-actin in regions where it is enriched (e.g. at the cell cortex) and thus indirectly promote microtubule destabilization. Of note, Maspin RCL motif was also predicted to interact with a region in the β-actin homodimer (aa 348-355) which overlaps with a common site for actin-binding proteins (Dominguez, 2004). In addition, a short sequence in Maspin (aa 110-116) is a putative G-actin-binding RPEL motif (Diring et al., 2019). How Maspin RCL, whose molecular function remains unknown, precisely coordinates the crosstalk between microfilaments and microtubules *in vitro* and in cells is a topic for future studies.

### Maspin depletion drives cells to an intermediate EMT state

In our proteomic assay, we failed to capture classical cell adhesion molecules such as E-cadherin or even β1-integrin, which is a known Maspin interacting protein (Cella et al., 2006). This may be due to our different experimental conditions or model system. Given that Maspin was observed at the cell cortex (Fig. 2A and B) and it interacts with actin filaments *in vitro* (figure 1I), we hypothesized that it may be part of the adhesion complexes which connect the microenvironment (extracellular matrix and adjacent cells) to the cytoskeleton. To substantiate this, we integrated our AP/MS-identified interacting proteins to an Adhesome dataset, manually curated from different large-scale proteomic studies (Fig. 5A). This Adhesome is mainly composed of hemidesmosomes (Integrin-mediated), adherens junctions and desmosomes (E-cadherin and Desmoplakin) and cortical cytoskeleton components (Suppl. Table 2). We found that Adhesome and Maspin share 11 interacting proteins (Fig. 5B). They were classified mainly as adhesion-cytoskeleton related terms by GO molecular function analysis (Fig. 5C). Interestingly, Maspin was repeatedly captured as part of different adhesion plaques in most of these studies, co-occurring with plakins such as Plectin (Fig. 5B), which was captured in our AP/MS analysis (Fig. 1E) and interacts with integrins (Koster et al., 2004). Indeed, Desmoplakin, another plakin family member, interacts with Plectin (Eger et al., 1997) and has been recently shown to interact with Maspin in HaCaT cells (Rathod et al., 2024). Plakins are mainly cytolinkers, connecting cell adhesion molecules to the cortical cytoskeleton (Suozzi et al., 2012; Liem, 2016; Zhang et al., 2017a). They maintain epithelial cell polarity, preventing EMT in MCF-10A cells (Geay et al., 2024). During EMT, cell shape undergoes modifications as a consequence of cell-cell contact disruption, cytoskeleton reorganization and transcriptional activation of mesenchymal genes (Thiery et al., 2009; Kalluri and Weinberg, 2009; Yang et al., 2020; Nurmagambetova et al., 2023). Since we noticed that Maspin KO cells partially lose their ability to arrange as regular packed clusters, occasionally exhibiting an elongated shape (Fig. 5D), we reasoned that our Maspin KO cells had undergone EMT. Indeed, most of the phenotypic changes observed for microfilaments and microtubules upon Maspin depletion resemble activation of an EMT program. For instance, Maspin KO cells display an increased size (Fig. 3E) and lose most of the regular cortical cytoskeleton distribution, assembling more actin stress fibers and reorganizing their microtubules (Fig. 3A-C) (Nurmagambetova et al., 2023). We also observed cytoplasmic vimentin dispersion upon Maspin transient knockdown (Suppl. Fig. 2B). In addition, EMT induction results in more dynamic microtubules, as recently measured by tracking EB3 in epithelial cells (Nurmagambetova et al., 2023). This is consistent with the increase of EB3 comets growth speed observed in our MCF- 10A KO cells (Fig. 4D).

**Figure 5.**
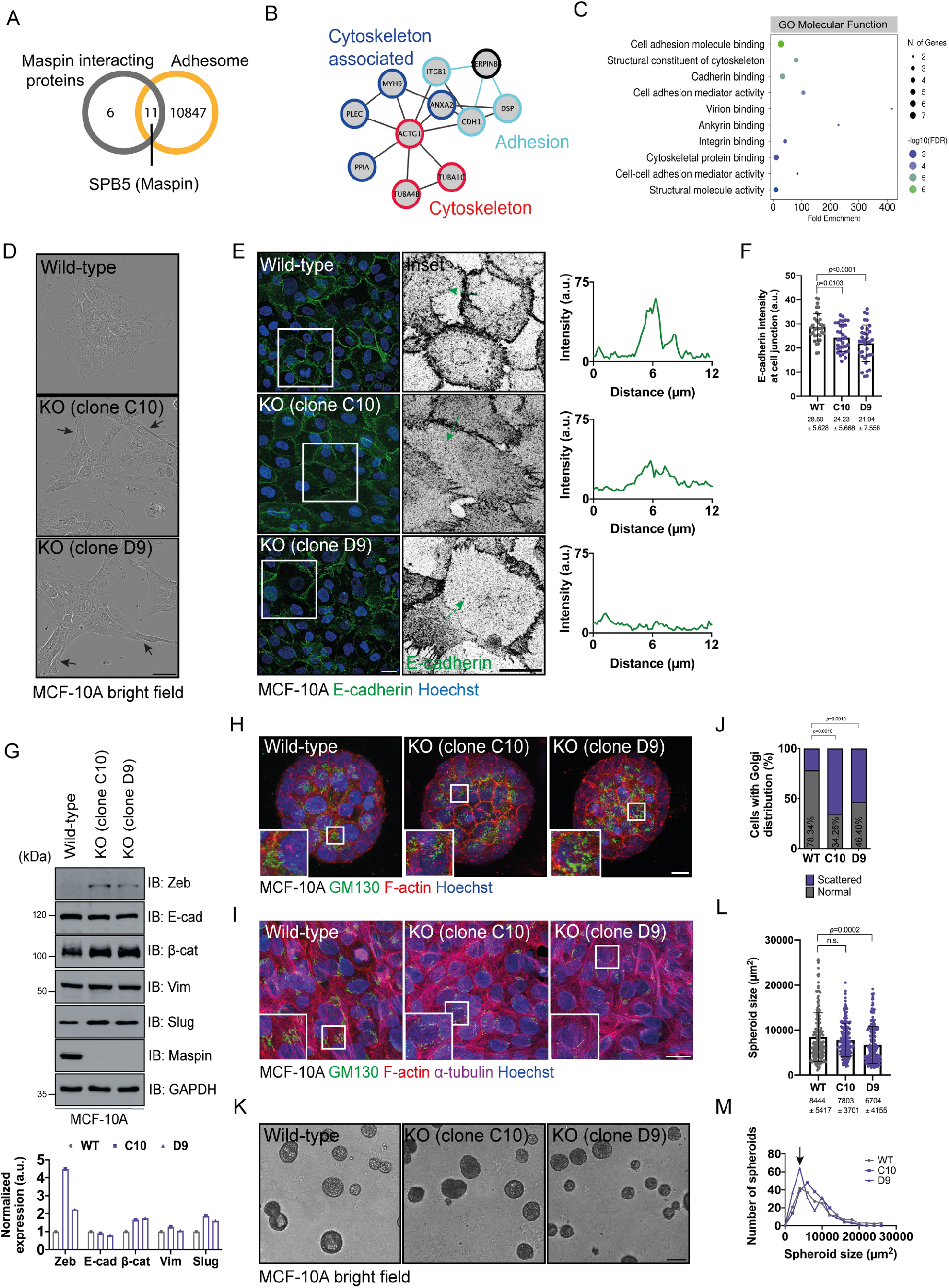
Maspin depletion triggers an intermediate EMT state. (A) Venn diagram showing the intersection between identified Maspin interacting proteins and the Adhesome dataset; **(B)** Interactome for shared interacting proteins (right) include adhesion - Desmoplakin (DSP), β1-integrin (ITGB1), E-cadherin (CDH1); cytoskeleton - Actin gamma (ACTG1), Tubulin (TUBA1C and TUBA4B); and cytoskeleton-associated proteins - Plectin (PLEC) and myosin heavy chain-9 (MYH9). Black edges represent physical interactions reported by String, whereas cyan edges represent interactions experimentally described by others (DSP (Rathod et al., 2024), ITGB1 (Cella et al., 2006) and CDH1 (Guo et al., 2014); **(C)** GO Molecular Function component showing the top-10 terms ranked by FDR score, including cell-cell, cell-substrate adhesion and cytoskeleton-related terms depicted in (B); **(D)** Bright-field images of MCF-10A WT cells and KO clones (C10 and D9). Black arrows indicate distinct spatial organization for KO clones (elongated) compared to WT cells (packed). Scale bar: 20 µm; (**E**) E-cadherin labeling in MCF-10A WT cells and KO clones. Scale bar: 10 µm; **(F)** E-cadherin fluorescence intensity at cell junction for MCF-10A WT (n = 36 cells), KO C10 (n = 33 cells) and KO D9 (n = 35 cells); **(G)** Western blot analysis for the EMT markers Zeb, E-cadherin, β-catenin, Vimentin and Slug in MCF-10A WT and KO cells. GAPDH was used as a loading control and Maspin knock-out probed with an anti-Maspin antibody. Expression levels were quantified by densitometry (lower graphic); **(H)** Representative immunofluorescence images of F-actin and GM130 in MCF-10A WT, KO C10 and KO D9 acini at day 12. Scale bar: 10 µm; **(I)** Representative immunofluorescence images of F-actin, GM130 and α-tubulin in MCF-10A WT, KO C10 and KO D9 cells. Scale bar: 15 µm; **(J)** Percentage of cells with abnormal (scattered) *versus* normal Golgi distribution, MCF-10A WT (n = 173 cells), C10 (n = 181 cells) and D9 (n = 155 cells); **(K)** Bright-field images of on-top tridimensional MCF-10A WT, KO C10 and KO D9 acini at day 12 after seeding. Scale bar. 100 µm; **(L)** Quantification of spheroid size for MCF-10A WT (n = 200 acini), KO C10 (n = 209 acini) and KO D9 (n = 212 acini) imaged in (K); **(M)** Frequency distribution of spheroids quantified in (L) according to size. Arrow indicates a higher frequency of smaller spheroids for MCF-10A KO D9 cells.

To confirm that loss of Maspin triggers an EMT state, we quantified protein levels of EMT markers, including E-cadherin, β-catenin, vimentin, Slug and Zeb. Notably, we found a mixed pattern for E-cadherin labeling in MCF-10A KO cells, which appeared either as spiky extensions resembling filopodia projections or as cytoplasmic puncta. On the other hand, MCF-10A WT cells exhibit continuous, linear E-cadherin, at cell-cell junctions (Fig. 5E). This is consistent with cell cortex changes promoted by actin remodeling to favor cell motility during EMT (Vasioukhin et al., 2000; Lamouille et al., 2014) and aligns with Maspin inhibition of cell motility, as previously reported (Sheng et al., 1996). In MCF-10A siMaspin cells, in which Maspin expression levels are not entirely downregulated, E-cadherin appears dotted at cell junctions (Suppl. Fig. 3A), an indication of cell-cell contact disruption. This becomes even more evident for EpH4 shMaspin cells (Suppl. Fig. 3C) and suggests that the Maspin-driven EMT state depends on its expression levels. Finally, E-cadherin intensity at the cell cortex partially decreased (Fig. 5F and Suppl. Fig. 3B and D) likely due to its mislocalization, but also due to a slight reduction of its expression levels, as indicated by Western blot (Fig. 5G). Concomitantly, we noticed the upregulation of mesenchymal markers such as β-catenin, Vimentin, Slug and Zeb (Fig. 5G). Some of these modifications such as increased Slug expression were confirmed in HaCaT KO cells as well (Suppl. Fig. 3E). These results agree with the altered E-cadherin basolateral pattern found in the prostate tissue of Maspin heterozygous mice (Shao et al., 2008).

So far, our phenotypic observations indicate that loss of Maspin triggers an EMT state in epithelial cells by acting at the interface between the cytoskeleton and adhesion sites. However, epithelial cells undergo several intermediate states before full conversion to mesenchymal cells (Kalluri and Weinberg, 2009). In order to better characterize our EMT state, we cultured MCF-10A cells on top of basement membrane-enriched matrigel substrate. These cells form acini-like spheroids under this particular condition, which allows better cell polarization, an important feature that is primarily altered during EMT. During this process, apical-basal polarity becomes predominant and the Golgi apparatus locates apically, towards the hollow lumen of each acinus (Debnath et al., 2003). Opposingly, epithelial cells undergoing an EMT program may display mispositioned Golgi and progressively reverse their polarity during mammary morphogenesis (Burute et al., 2017). To evaluate if loss of Maspin results in Golgi reorganization and polarity reversal, we cultured MCF-10A WT and KO spheroids for 12 days and co-stained them for Golgi (GM130) and F-actin. Interestingly, KO cells exhibit more fragmented Golgi (Fig. 5H), which we confirmed in monolayer cell culture (Fig. 5I and J). This may indicate that the cytoskeleton rearrangements upon Maspin depletion disrupt Golgi spatial arrangement, possibly resulting from microtubule reorganization in Maspin KO cells. Based on the observations described above and on the lack of full polarity reversal in KO cells, we conclude that Maspin loss leads to an intermediate EMT state in our model system. This result is in line with previous data suggesting that Maspin deficiency in mice induces polarity reversal, as observed by mislocalized apical junction marker ZO-1 in prostate tissue (Shao et al., 2008). In addition to these observations, Maspin KO acini exhibit a reduced size on average (Fig. 5K and L), with a higher frequency of smaller spheroids observed for KO clone D9 (Fig. 5M). We presently do not understand how this observation relates to Maspin-cytoskeleton interaction. We conclude that Maspin either promotes or suppresses tumor progression by modulating an EMT hybrid phenotype. This balance likely depends on cell and tissue context, as well as on its subcellular localization, as previously proposed (Nosaka et al., 2015; Goulet et al., 2012; Longhi et al., 2021).

## Materials and methods

### Cell culture and treatments

Immortalized human mammary epithelial cells (MCF-10A) (BCRJ - Banco de Células do Rio de Janeiro) were cultured in DMEM/F12 (12500062, Gibco) containing 5% donor horse serum (26050088, Gibco), 20 ng/mL EGF (PHG0311, Thermo Fisher Scientific), 0.5 mg/mL hydrocortisone (H0888, Sigma-Aldrich), 100 ng/mL cholera toxin (C8052, Sigma-Aldrich), 10 µg/mL insulin (I0516, Sigma-Aldrich) and 1x penicillin-streptomycin (15070063, Gibco). Immortalized mouse mammary epithelial cells (EpH4) (a kind gift from Dr. Alexandre Bruni-Cardoso, Department of Biochemistry, University of Sao Paulo) were cultured in DMEM/F12 supplemented with 5 µg/mL insulin, 50 µg/mL gentamicin (15750078, Gibco), 1 µg/mL puromycin (P9620, Sigma-Aldrich) and 2% fetal bovine serum (Cultilab). Immortalized human keratinocyte cells (HaCaT) (BCRJ - Banco de Células do Rio de Janeiro) were cultured in DMEM (12100061, Gibco) supplemented with 10% fetal bovine serum and 1x penicillin-streptomycin. All cell lines were kept at 37 °C and 5% CO2 in a humidified incubator (Thermo Fisher Scientific) and routinely trypsinized in 0.05% Trypsin-EDTA (25300054, Gibco).

Experiments with nocodazole: in order to enrich for mitotic cells (G2/M arrest and release), cells were treated with 50 ng/mL of nocodazole (sc-3518, Santa Cruz Biotech.) for 16 h, followed by two washes with warm PBS and release into fresh medium without nocodazole for 30 min. For microtubule regrowth assays, cells were kept in a complete medium containing 1 µg/mL of nocodazole for 1 h, then washed twice with warm PBS and recovered in drug-free fresh medium at 37 °C.

### RNAi-mediated knockdowns

For transient knockdown, MCF-10A cells were cultured in complete medium in a p6-multiwell plate for 24 h. For siRNA transfection, 150 µL of Opti-MEM I (31985062, Gibco) and 8 µL of Lipofectamine RNAiMAX (13778150, Invitrogen) were added to tubes. siRNAs (120 pmol, stock solution 20 nmol) were added on top of each individual tube and incubated for 10 min at room temperature. Cell medium was changed shortly before transfection and final mixtures were added dropwise and cells were incubated for 72 h. The following commercially available siRNA sequences were used: Stealth siRNAs targeting Maspin mRNA sequence - HSS1079<u>60</u> (siMaspin #60) and HSS1822<u>95</u> (siMaspin #95) (Thermo Fisher Scientific) - along with the negative control Stealth <u>N</u>egative <u>C</u>ontrol Low GC duplex (siNC) (452002, Thermo Fisher Scientific).,

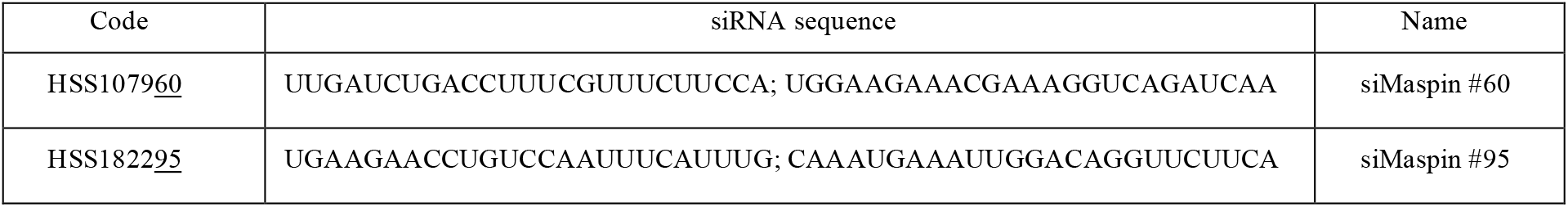

For stable knockdown, EpH4 cells were cultured in complete medium in a p24-multiwell plate 24h prior to transductions at 70% confluency. Next, fresh medium containing 8 µg/mL of polybrene (TR-1003-G, Sigma-Aldrich) was added to each well and commercially available lentiviral particles (SHCLNV-NM_009257, MISSION^®^ Lentiviral Transduction Particles, Sigma-Aldrich) containing the lentiviral vector pLKO.1-puro (SHC001, Sigma-Aldrich) were added dropwise on top of cells at a MOI (Multiplicity of infection) value of 1.3. After 24 h, medium was changed and cells allowed to recover for an additional 24 h. Next, puromycin selection (2.5 µg/mL) was carried out for 2 weeks and cells were further cultured in the presence of 1.0 µg/mL of puromycin. The following shRNA sequences targeting Maspin mRNA along with the negative control (SHC001V, Sigma-Aldrich) were used:

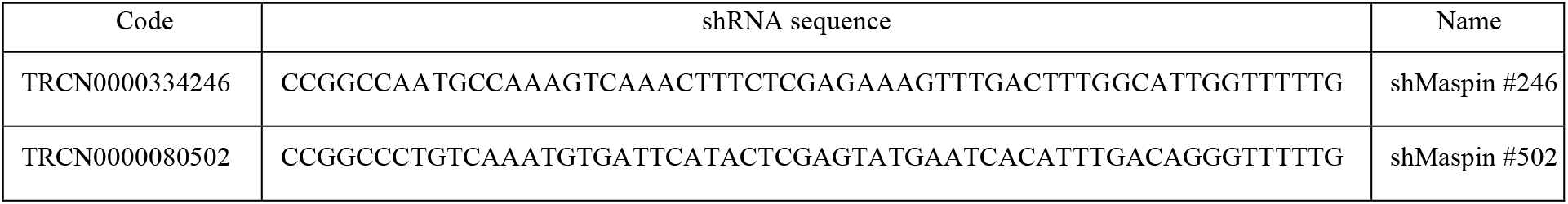

All Maspin knockdowns were confirmed by Western blot and immunofluorescence with anti-Maspin antibodies.

### Generation of Maspin knock-in and knock-out cells by CRISPR/Cas9

CRISPR/Cas9 knock-in was performed using Cas9 Nuclease 2-NLS (Synthego), sgRNAs targeting Maspin gene sequence (N1/N2-terminal and C-terminal) and a template DNA generated by PCR containing a GFP and Puromycin resistance sequence (A42992, Invitrogen). The following primers were used:

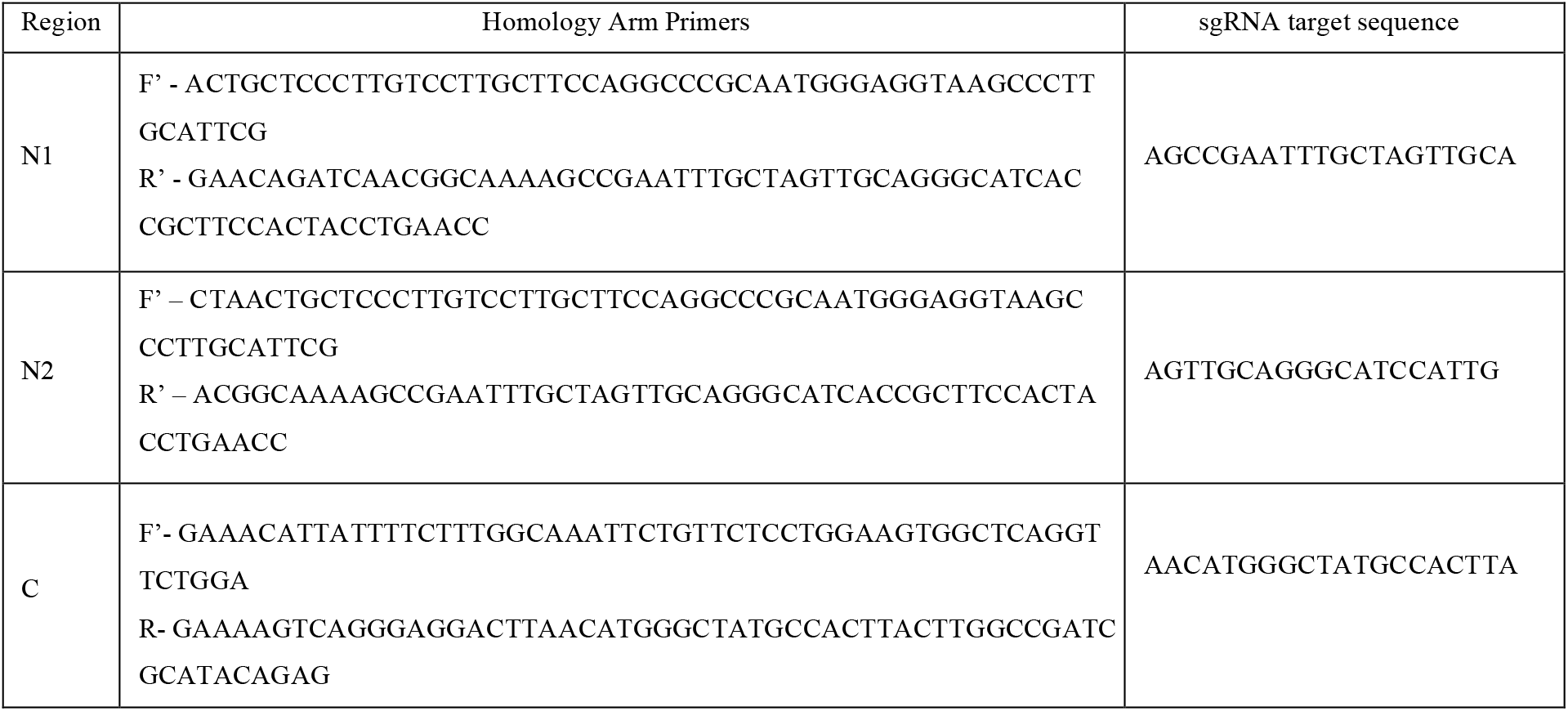

PCR mixes containing 1 U of Q5 polymerase (M0491S, New England Biolabs), 1x reaction buffer, 10 mM dNTPs, 0.5 µL of each template (GFP-Puro), 10 µM primers and nuclease free H2O (up to 50 µL) were ran on a DNA Engine TETRAD2 Peltier Thermal cycler (MJ 426 Research) with the following program: initial denaturation at 98°C for 10 min followed by 30 cycles of denaturation at 98 °C for 30 sec, annealing at 56 °C for 25 sec and elongation at 72 °C for 1 min, final elongation at 72 °C for 7 min and a 4 °C hold. PCR products were visualized on a 1 % agarose gel (ChemiDoc MP Imaging System, Bio-Rad) and the amplicons were further purified (T1030L, New England Biolabs) prior to mixing with RNP complexes.

CRISPR/Cas9 knock-out was performed using Cas9 Nuclease 2-NLS (Synthego) and a mix of 3 different sgRNAs (Multi-guide system, Synthego) targeting exon 2 of Maspin with the following sequences: sgRNA1 (AAGCCGAATTTGCTAGTTGC), sgRNA2 (GAGAAGAGGACATTGCCCAG) and sgRNA3 (CACTGCAAATGAAATTGGAC).

For both knock-in and knock-out, RNP complexes were formed in Nucleofector solution at a 6:1 ratio (sgRNA:Cas9) and immediately transfected by Nucleofection into MCF-10A or HaCaT cells using the Amaxa SF Cell Line 4D-Nucleofector X kit S (Lonza, PBC2-00675) and the program DS-138 (MCF-10A and HaCaT) on the 4D-Nucleofector X unit. Cells were harvested and 4 x 10^5^ cells were added to the RNP complex mix in a 16-well Nucleocuvette. After nucleofection, the transfected cells were resuspended in conditioned medium DMEM/F12 (MCF-10A) or DMEM (HaCaT), plated in p12- multiwell plates, and 24 h later cells were fed with conditioned media. This conditioned media was previously obtained by pooling and filtering the supernatants of MCF-10A or HaCaT confluent cells after 72 h and 96 h of subculturing, and by mixing them with fresh complete medium at a 1:1 ratio.

After nucleofection, knock-in cells were cultured in conditioned media, allowed to reach 70% confluency and split into three wells of a p12-multiwell plate. Puromycin selection was carried out after 24 h at different doses (0.5 µg/mL; 0.75 µg/mL and 1µg/mL) for two weeks and fluorescence signal monitored using Incucyte (Sartorius). MCF-10A knock-ins were then scaled up and routinely cultured in the presence of puromycin (0.5 µg/mL for C-terminal tagged; 1 µg/mL for N1/N2-terminal tagged).

After nucleofection, knock-out cells were allowed to reach 90% confluency and the edited pool was sequenced to confirm knock-out efficiency. Single-cell cloning was then performed using DispenCell Single-Cell Dispenser (Molecular Devices) according to manufacturer’s instructions and cells were individually seeded in a 96-multiwell plate containing conditioned media at 37 °C. The plate was monitored weekly for colony formation until 80% confluency was reached (around 2-weeks). Single cell colonies were then expanded in 24-multiwell plates containing fresh media and further scaled up. Genomic DNA was extracted (T3010L, New England Biolabs) from at least 5 different clones and PCR was conducted with primers targeting a region on exon 2 for human Maspin (F’ - ACAGAAAGTTCCTTGTTTCTTCTGA; R’ - GCCTGTCCCAGATGTGAGTT). PCR mixes containing 1 U of PfuX7 polymerase, 1x reaction buffer, 10 mM dNTPs, 50 ng of genomic DNA, 10 µM primers and nuclease free H2O (up to 50 µL) were ran on a DNA Engine TETRAD2 Peltier Thermal cycler (MJ Research) with the following program: initial denaturation at 98 °C for 10 min followed by 30 cycles of denaturation at 98 °C for 30 sec, annealing at 54 °C for 25 sec and elongation at 72 °C for 1 min, final elongation at 72°C for 7 min and a 4 °C hold. PCR products were visualized on a 1% agarose gel (ChemiDoc MP Imaging System, Bio-Rad), purified (T1030L, New England Biolabs) and sent for Sanger Sequencing (Génome Québec) with a nested sequencing primer (TTCCTTGTTTCTTCTGAGATCAATTACTTT). Sequencing data was analyzed with Synthego’s ICE software, available at: https://www.synthego.com/products/bioinformatics/crispr-analysis.

### Molecular cloning, protein expression and purification

The coding sequence for full-length human Maspin protein was PCR amplified from previously described GFP-Maspin construct (Reina et al., 2019) using PfuX7 polymerase and the following primers: (F’GGGGAGCUATGGATGCCCTGCAACTAG;R’GGGGAACUAGGAGAACAGAATTTGCCAAAG). This amplicon was inserted into a modified pHAT-HGUS vector containing an N-terminal 6xHis-tag followed by EGFP, and a C-terminal Strep-tag II using the USER enzyme cloning system (Geu-Flores et al., 2007).

For His-GFP-Maspin-StrepII protein, we used a similar protein expression protocol previously described (McAlear and Bechstedt, 2022). *E. coli* BL21 (DE3) containing protein expression vectors were grown to O.D. 0.6 at 37 °C, and expression was induced using 0.5 mM Isopropyl β-D-1- thiogalactopyranoside (IPTG) at 18 °C for 16 h. Bacterial pellets were harvested by centrifugation and resuspended in Buffer A (50 mM Na2HPO4, 300 mM NaCl, 4 mM imidazole, pH 7.8). Cells were lysed using a French press (EmulsiFlex-C5, Avestin). Constructs containing both His and Strep tags were purified using gravity flow columns containing His60 Ni-NTA resin (Clontech) followed by Streptactin affinity chromatography (IBA Lifesciences, Germany). Purified Maspin was eluted with BRB80 (80 mM PIPES-KOH, pH 6.85, 1 mM EGTA, 1 mM MgCl2) or Tris-HCl (100 mM Tris-HCl pH 7.5, 150 mM NaCl, 1 mM EDTA) containing 2.5 mM desthiobiotin and 10% glycerol. Purified Maspin was always used fresh for each experiment. Protein purity was analyzed by SDS-PAGE gel stained with Coomassie blue. Protein concentration was determined by absorbance at 510 nm with a DS-11 FX spectrophotometer (DeNovix, Inc) (McAlear and Bechstedt, 2022).

Tubulin was purified from bovine brains as previously described (Ashford and Hyman, 2006) with the modification of using Fractogel EMD SO3- (M) resin (Sigma-Aldrich) instead of phosphocellulose. Tubulin was labeled using Atto-633 NHS-Ester (ATTO-TEC) and tetramethylrhodamine (TAMRA, Invitrogen) as described (Hyman et al., 1991). An additional cycle of polymerization/depolymerization was performed before use. Protein concentrations were determined using a DS-11 FX spectrophotometer (DeNovix, Inc).

The pGEX-4T1 and pGEX-Maspin constructs were previously described (Cella et al., 2006) and used to express control GST-only and GST-Maspin bait proteins, respectively. *E. coli* BL21(DE3) containing protein expression vectors were grown to O.D. 0.6 at 37 °C, and expression was induced using 0.8 mM IPTG at 37 °C for 4 h. Bacterial pellets were harvested by centrifugation and resuspended in Buffer T1 (50 mM Tris-HCl, pH 7.5, 150 mM NaCl, 1% Triton X-100, 5 mM MgCl2, 1 mM PMSF, and 10 µg/mL leupeptin and aprotinin). Cells were lysed on ice by sonication (10 cycles of 30 sec each with intervals of 1 min, 70% amplitude) (Vibra-Cell VC 505 sonicator (SONICS, Newtown, CT, USA). The material was centrifuged at 14,000 rpm at 4 °C for 30 min, and the soluble fractions containing the GST-only or GST-Maspin proteins were collected, frozen at -80 °C and used for further affinity purification experiments (da Silva et al., 2020). An SDS-PAGE followed by Coomassie Brilliant Blue R-250 (Bio-Rad, Hercules, CA, USA) staining was used to assess protein purity after affinity purification. GST-fusion protein concentration was estimated based on a standard curve using bovine serum albumin (BSA) at different known concentrations in the same gel. This gel was scanned with an Odyssey Imager (LI-COR, Lincoln, NE, USA) operated with dedicated Odyssey V 3.0 software (LI-COR, Lincoln, NE, USA), which was used for band quantification in ImageJ.

### Pull-down and Western blot

For pull-down experiments, the bacterial lysates were separately incubated with glutathione agarose (16100, Thermo Fisher Scientific) and left rotating for 1.5 h at 4 °C. The beads were washed 6x with buffer T2 (50 mM Tris pH 7.5, 0.5% Triton X-100, 150 mM NaCl, 5 mM MgCl2 and 1x protease inhibitor cocktail) with intercalated centrifugations at 800 x *g* for 3 min at 4 °C. Supernatants from the final washes were removed and freshly harvested MCF-10A cell lysates were added to the pellets. The reaction mixtures were allowed to rotate at 4 °C for 1 h and then centrifuged at 800 x *g* for 5 min at 4 °C. The beads were washed three times with buffer I (Tris 25 mM pH 7.3, 150 mM NaCl) and eluted with buffer G (reduced glutathione 10 mM, 150 mM NaCl, 1 mM DTT, 50 mM Tris-HCl pH 8.5) under constant rotation at 4 °C. An aliquot of 5 µL of final supernatants were run on an SDS-PAGE gel and silver staining carried out to confirm recombinant protein elution. For pull-down validations, 10 µL of each eluate were denatured in 3x Laemmli buffer and immunoblot performed with specific primary antibodies. MCF-10A cell lysates were obtained by scraping cells in modified buffer I (Tris 25 mM pH 7.3, 150 mM NaCl, 0.1% IGEPAL, 1x protease cocktail inhibitor, 1 mM NaF, 1 mM Na3VO4) and thoroughly pipetting to avoid aggregates. Then, a sonication step was done on ice (3 cycles, 7 sec each with 30 sec intervals and 20% amplitude) followed by a final centrifugation at 14.000 rpm 4 °C for 10 min. The obtained supernatants were freshly used for incubation with recombinant proteins on the same day they were purified.

Western blot analysis was performed as previously described (Longhi et al., 2021) and quantified by densitometry using ImageJ. Primary antibodies were diluted in TBS-T only: mouse anti- Maspin (1:2000, clone 5C6.2, MABC603, Millipore); mouse anti-GAPDH (1:1000, GA1R, MA5- 15738, Invitrogen), mouse anti-α-tubulin (1:1000, clone B7, sc-5286, Santa Cruz Biotech.), mouse anti-E-cadherin (1:1000, HECD-1, 13-1700, Invitrogen), mouse anti-β-catenin (1:1000, sc-7963 Santa Cruz Biotech.), mouse anti-Vimentin (1:1000, #CBL202, Chemicon), mouse anti-Slug (1:1000, A-7, sc- 166476, Santa Cruz Biotech.), mouse anti-Zeb1 (1:1000, H-3, sc-515797, Santa Cruz Biotech.), mouse anti-GFP (1:1000, GF28R, MA5-15256, Invitrogen). Secondary antibodies were diluted in skim milk 5% in TBS-T: goat anti-mouse HRP (1:10.000, 1721011 Bio-Rad).

### Proteomics, gene ontology and interactomes

The samples were digested and cleaned as previously described (De Jesus et al., 2016). Briefly, eluates were resuspended in 400 µL of ice-cold 100 mM Tris-HCl buffer at pH 8.5. For protein precipitation, 100 µL of cold trichloroacetic acid (TCA) was added and samples were incubated overnight at 4°C. After centrifugation at 14,000 rpm for 30 min at 4°C, pellets were washed twice with ice-cold acetone and allowed to dry at room temperature. The protein concentration in each sample was measured using the Pierce BCA Protein Assay Kit (Thermo Fisher Scientific) following manufacturer’s recommendations. For protein digestion, 100 µg of protein were resuspended in 30 µL of a solution containing 8 M urea, 75 mM NaCl, and 50 mM Tris at pH 8.2. Proteins were reduced with 5 mM DTT for 30 min at room temperature, followed by alkylation with 14 mM iodoacetamide in the dark for 30 min. Subsequently, protein extracts were digested overnight with 1 µg of trypsin (Promega, Madison, Wisconsin) in a solution containing 10 mM Tris-HCl at pH 8.0 and 2 mM CaCl2, maintained at 37°C.

The digestion reactions were halted by the addition of 5% formic acid, and the samples were vacuum dried. After protein digestion, the samples were redissolved in 400 µL of 0.1% TFA and desalted using the Sep-Pak Light tC18 column (Waters, Milford, Massachusetts). The desalted samples were then dried and prepared for analysis by mass spectrometry.

Peptides were resuspended in 0.1% formic acid and injected into a custom-made 5 cm reversed-phase pre-column (inner diameter 100 µm, filled with 10 µm C18 Jupiter resins - Phenomenex, Torrance, California) connected to a nano HPLC system (NanoLC-1DPlus, Proxeon, Thermo Fisher Scientific, Waltham, Massachusetts). Peptide fractionation was performed on a custom 10 cm reversed-phase capillary emitter column (inner diameter 75 µm, filled with 5 µm C18 Aqua resins - Phenomenex) using a gradient of 3–35% acetonitrile in 0.1% formic acid over 45 min, followed by a gradient of 35– 95% for 5 min, maintained at 95% for 8 min and, 95–5% for 2 min at a flow rate of 300 nL/min. The eluted peptides were directly analyzed using an LTQ-Orbitrap Velos mass spectrometer (Thermo Fisher Scientific). The source voltage and capillary temperature were set to 1.9 kV and 200 °C, respectively. The mass spectrometer operated in a data-dependent acquisition mode, automatically switching between one Orbitrap full scan and ten ion trap tandem mass spectra. FT scans were acquired from m/z 200 to 2000 with a mass resolution of 30,000. MS/MS spectra were obtained at a normalized collision energy of 35%. Singly charged and charge-unassigned precursor ions were excluded, with dynamic exclusion set to 45 sec for exclusion duration, 500 for exclusion list size, and 30 sec for repeat duration.

Raw data were processed using MaxQuant software version 1.6.10.43 and the Andromeda search engine. Proteins were identified by searching against the complete database sequence of *Homo sapiens* (downloaded from UniProt - UP000005640- 82,518 entries) along with a set of commonly observed contaminants. Carbamidomethylation (C) was set as a fixed modification, while oxidation (M) and acetylation (N-terminal) were set as variable modifications. The maximum number of modifications per peptide was set to 5, with a maximum of 2 missed cleavages. The MS1 tolerance was 6 ppm and the MS2 tolerance was 0.5 Da. The maximum false discovery rates for peptides and proteins were set to 0.01. For matching between runs, a time window of 2 min was used. Bioinformatics analysis was performed using Perseus software (http://www.perseus-framework.org/). Proteins matching the reverse (or contaminants) database, or identified only by modified peptides, were filtered out. Proteins were identified by at least one unique peptide. Relative protein quantification was carried out using the LFQ algorithm of MaxQuant, with a minimum ratio count of two. Protein quantification was based on LFQ values of "razor and unique peptides" of unmodified peptides, using the Proteingroups.txt. We kept proteins exclusively found or enriched (LFQ > 1.5-fold) in at least 2 biological replicates from GST- Maspin baits compared to GST-only baits. An average for LFQ values corresponding to each protein hit was taken and a final normalization factor was set for all hits not found based on the lowest LFQ value found for a hit within each group (GST- or GST-Maspin). Fold change was then calculated by taking the mean values of GST-Maspin hits and dividing them by the corresponding mean values of GST-only control hits (Suppl. Table 1). A final heatmap was plotted with the adjusted fold change values using GraphPad Prism 8.0.

GO enrichment analysis was performed by combining proteins identified in DMSO-treated samples and Nocodazole-treated samples (gene list) and submitting them to the Shiny GO 0.80 platform (Ge et al., 2020). An FDR of 0.05 was set as a cutoff to select the top 10 most significant molecular functions (*Homo sapiens*). Interactomes were generated using String database (*Homo sapiens*), medium confidence (0.400) and FDR of 0.05. For the Adhesome dataset, protein lists from each study were downloaded and manually organized (Suppl. Table 2). Venn diagrams were generated by intersection of protein hits using https://bioinformatics.psb.ugent.be/webtools/Venn/. All networks were imported into Cytoscape and nodes and edges customized according to protein classes and direct physical interactions, respectively. The latter was complemented by manual annotations of previous studies where interactions were found, but not reported by the String database.

### In silico protein-protein interaction prediction

Protein complexes were generated by inputting the human sequences of SerpinB5 (P36952), TBA1A (Q71U36), TBB1 (Q9H4B7) and ACTB (P60709) into AlphaFold (Bryant et al., 2022). Complexes were colored and analyzed using ChimeraX v. 1.8.

### F-actin binding assay

*In vitro* binding of Maspin to F-actin was performed using rabbit skeletal muscle actin (Cytoskeleton, #AKL99). Briefly, purified actin was reconstituted in General Actin buffer (5 mM Tris-HCl pH 8.0, 0.2 mM CaCl2) according to manufacturer’s instructions. Actin was incubated on ice for 30 min and centrifuged at 14,000 rpm for 15 min at 4 °C. The supernatant (200 µL) was transferred to a new tube and 20 µL of Actin Polymerization buffer (500 mM KCl, 20 mM MgCl2, 10 mM ATP) were added along with 1 µL of labeled actin. The reaction mixture was incubated at 30 °C for 30 min and centrifuged at 100,000 x *g* in an Airfuge (Beckman-Coulter) for 10 min. For TIRF experiments, flow channels were first incubated with a polyclonal rabbit anti-actin antibody (Biomedical Technologies Inc., BT-560) diluted in 1x KMEI buffer (50 mM KCl, 1 mM MgCl2, 1 mM EGTA, 10 mM imidazole pH 7.0) at 1:50 and then blocked with 5% Pluronic F-127. Channels were washed twice with BRB80 and polymerized F-actin diluted in TIRF imaging buffer (1x KMEI, 100 mM DTT, 0.2 mM ATP, 15 mM glucose, 0.5% methyl cellulose, 0.01 mg/mL catalase, 0.05 mg/mL glucose oxidase, 0.1% BSA) and loaded into the channels (Jiang and Huang, 2017). Images were taken prior to flowing in recombinant Maspin through the channel.

### Paclitaxel-stabilized microtubules and dynamic microtubule assays

For paclitaxel-stabilized microtubules, a polymerization mixture was prepared with BRB80 + 32 µM tubulin + 1 mM GTP + 4 mM MgCl2 + 5% DMSO. The mixture was incubated on ice for 5 min, followed by incubation at 37 °C for 30 min. The polymerized microtubules were diluted into prewarmed BRB80 + 10 µM paclitaxel, centrifuged at 110,000 rpm (199,000 × g) in an Airfuge (Beckman-Coulter), and resuspended in BRB80 + 10 µM paclitaxel (99.5+%, GoldBio).

To visualize dynamic microtubules, we reconstituted microtubule growth off of GMPCPP double stabilized microtubule ‘seeds’ (Gell et al., 2010). In short, cover glass was cleaned in acetone, sonicated in 50% methanol, sonicated in 0.5 M KOH, exposed to air plasma (Plasma Etch) for 3 min, then silanized by soaking in 0.2% dichlorodimethylsilane in n-heptane. 5 µl flow channels were constructed using two pieces of silanized cover glasses (22 × 22 and 18 × 18 mm) held together with double-sided tape and mounted into custom-machined coverslip holders. GMPCPP seeds were prepared by polymerizing a 1:4 molar ratio of TAMRA labeled:unlabeled tubulin in the presence of GMPCPP (Jena Biosciences) in two cycles, as described previously (Gell et al., 2010). Channels were first incubated with anti-TAMRA antibodies (Invitrogen) and then blocked with 5% Pluronic F-127. Flow channels were washed three times with BRB80 before incubating with GMPCPP seeds. On each day of experiments, tubes of unlabeled and Atto-633 labeled tubulin were thawed and mixed at a 1:17 molar ratio,subaliquoted and refrozen in liquid nitrogen. For consistency in microtubule growth dynamics, one subaliquot of tubulin was used for each experiment. Microtubule growth from GMPCPP seeds was achieved by incubating flow channels with 8 µM tubulin in imaging buffer: BRB80, 1 mM GTP, 0.1 mg/mL bovine serum albumin (BSA), 10 mM dithiothreitol, 250 nM glucose oxidase, 64 nM catalase, and 40 mM D-glucose.

### Live cell imaging

Cells were seeded in a p12-multiwell plate and monitored using the Incucyte Live-Cell Analysis System (Sartorius) in a humidified incubator at 5% CO2 and 37 °C. Acquisition time was set at 5 min time intervals for asynchronous cell population or at 2 h time interval for knock-in cells over a total period of 24 h. At least 25 different fields per well were imaged using a 20x objective (bright field or 488 nm wavelength for knock-in cells).

### Immunofluorescence

Cells were cultured on top of glass coverslips previously covered with Poly-D-Lysine (1 mg/mL for MCF-10A, P6407, Sigma-Aldrich and 100 µg/mL for EpH4, A3890401, Gibco). Cells were fixed with PFA 4% in PBS for 10 min, washed twice with PBS and permeabilized with 0.5% Triton X-100 in PBS for 10 min at room temperature. Next, a blocking solution of 10% goat serum (G9023, Sigma-Aldrich) in PBS was added for 1 h at room temperature. Primary antibodies were diluted in blocking buffer and incubated overnight at 4 °C in a humidified chamber. Secondary antibodies were diluted in PBS and incubated for 1 h at room temperature in the dark. Nuclei were stained with Hoechst diluted in PBS (1:15,000) for 5 min. Glass coverslips were washed with PBS (10 min) and mounted with FluorSave (345789, Millipore) onto glass slides. Primary antibodies diluted in blocking solution were: mouse anti-α-tubulin (1:100, clone B7, sc-5286, Santa Cruz Biotech.), mouse anti-E-cadherin (1:100, HECD-1, 13-1700, Invitrogen), mouse anti-Maspin (1:100, clone 1F7, H00005268-M01, Abnova), rabbit anti-TUBA4A (1:100, SAB2102603, Sigma-Aldrich), mouse anti-GM130 (1:100, 610822, BD Biosciences). Secondary antibodies diluted in PBS were: goat anti-mouse Alexa Fluor 488 (1:500, A28175, Invitrogen), goat anti-rabbit Alexa Fluor 647 (1:500, A32733, Invitrogen). Rhodamine Phalloidin (R415, Invitrogen) was incubated along with secondary antibodies at 1:500 dilution.

### Image acquisition, quantification and statistical analysis

Data were acquired with a customized Zeiss Axio Observer seven equipped with a Laser TIRF III and 405/488/561/757-760 nm lasers, Alpha Plan-Apo 100 ×/1.46 Oil DIC M27, and Objective Heater 25.5/33 S1. Images on both systems were recorded on a Prime 95B CMOS camera (Photometrics) with a pixel size of 107 nm. For *in vitro* microtubule dynamics assay, images were taken at 10 sec time intervals for at least 16 min. For EB3 comets tracking, cells were seeded at 60% confluency in a 35 mm glass bottom dish and transiently transfected with 2000 ng of EB3-dtStayGold construct for 48 h (Hirano et al., 2022) using Lipofectamine 3000 according to manufacturer’s instructions. Cells were imaged at 500 msec time intervals for 2 min and microscope control was executed using ZenBlue (Zeiss).

Confocal images were acquired on an inverted Leica TCS SP8 confocal microscope equipped with a HC PL APO CS2 100x/1.4 NA objective lens. Confocal z-stack sections of 0.3-0.6 µm were acquired for 2D experiments and of 1.2 µm for 3D experiments. Image sets were acquired under the same gain and exposure time conditions within each experiment. Microscope control was executed using LasX software (Leica Microsystems).

All images were processed and analyzed using Fiji (ImageJ). EB3 comets were analyzed using kymographs. These were generated using KymographBuilder plug-in (ImageJ) and microtubule growth rates measured by manually drawing lines on kymographs and measuring the slope of growth using the Velocity Measurement Tool plug-in (ImageJ). The number of comets per area was counted using TrackMate (ImageJ) (Tinevez et al., 2017) based on the maximal intensity projection for the first three sequential frames recorded and dividing by cell area. The average anisotropy was measured using the FibrilTool plug-in (ImageJ) (Boudaoud et al., 2014). To this end, at least three regions of interest (defined ROI size) per cell were randomly selected and averaged based on EB3 maximal intensity projections. For microtubule regrowth assays, fluorescence intensity was measured based on a fixed circular ROI drawn and equally set for all MTOCs. Fluorescence intensity at cell junctions was measured manually drawing a 5-width segmented line around cell boundaries.

All statistical analyses and data plotting were performed using GraphPad Prism 8 (www.graphpad.com). In the scatter dot plots bars, dots represent independent measurements. Mean and standard deviation were represented below each graph. For numerical data, Shapiro–Wilk normality tests were applied and unpaired two-tailed Student’s t-tests were used where two independent groups were analyzed. For small sample size measurements (n < 25 cells), Mann–Whitney U test was employed instead. Multiple comparisons were done using One-Way ANOVA followed by Dunnett’s Multiple Comparison test. A *p*-value less than or equal to 0.05 was considered statistically significant. Figures were assembled using Adobe Illustrator and illustrations were assembled using BioRender.

## Online supplemental material

**Supplementary Table 1. List of all Maspin-binding proteins identified by proteomic analysis.** Maspin interacts with cytoskeleton components. Protein hits were eluted from GST-Pull-downs using Nocodazole-arrested or DMSO-treated MCF-10A cells.

**Supplementary Table 2. Adhesome dataset.** Intersection of Maspin-binding proteins with publicly available large-scale proteomic studies related to cell adhesion and cortical cytoskeleton components. **Video 1. Subcellular localization of endogenously tagged-Maspin in MCF-10A cells.** Maspin localizes to the leading edge of spreading cells. As confluency increases, it becomes diffuse in the cytoplasm of cells. Fluorescence signal was monitored by live imaging for 24 h (GFP-N1-Maspin knock-in).

**Video 2. Morphological changes observed for MCF-10A Maspin knock-out clones (C10 and D9) upon mitotic entry**. Loss of Maspin induces abnormal retraction tails in mitotic cells. Bright-field images were taken every 5 min over a total period of 5 h.

**Video 3. Visualization of EB3 comets in MCF-10A Maspin knock-out clones (C10 and D9)**. Increased microtubule growth speed observed for Maspin knock-out cells. Cells were transiently transfected with EB3-StayGold and imaged at 0.500 msec intervals. Video is accelerated to 10 frames per second.

**Video 4. TIRF-based *in vitro* reconstitution assay of dynamic microtubules.** Maspin suppresses microtubule growth. Atto633-tubulin and GFP-Maspin (50 nM or 500 nM) were added to immobilized GMPCPP-seeds and images were taken every 10 sec for at least 16 min.

## Author contributions

(LES) – Conceptualization, investigation, data curation, formal analysis, methodology, visualization, writing – original draft, writing – review and editing.

(LMGP) – Methodology, supervision, writing – review and editing. (APJM) – Investigation, formal analysis, validation, visualization.

(JPCC) – Conceptualization, funding acquisition, methodology, formal analysis, validation, visualization.

(SB) – Conceptualization, funding acquisition, investigation, methodology, supervision, formal analysis, validation, visualization, writing – review and editing.

(NC) – Conceptualization, funding acquisition, methodology, project administration, supervision, writing - original draft, writing – review and editing.

## Data availability

All data are available in the published article and its online supplemental material. The mass spectrometry proteomics data underlying Fig. 1 and Suppl. Table 1 have been deposited to the ProteomeXchange Consortium via the PRIDE repository with provisory reference number: 1-20240816- 114910-470199.

## Acknowledgments

We would like to thank Research Assistant Tatiana Lyalina (MSc.) from the Bechstedt laboratory for helping on molecular cloning, recombinant protein expression and purification and for assistance in all TIRF experiments performed in this study. For assistance with imaging using the confocal microscopes, we thank to Mr. Waldir Caldeira (MSc.) (Center for Image Acquisition and Microscopy at the Institute of Biosciences) and to laboratory specialist Adriana Yamaguti Matsukuma (MSc.) (Analytical Center at the Institute of Chemistry) of the University of São Paulo. Finally, we thank Dr. Vanessa Morais Freitas (Institute of Biomedical Sciences/University of Sao Paulo, Brazil) for contributing with reagents and equipment.

## Supplementary material

**Supplementary Figure 1.**
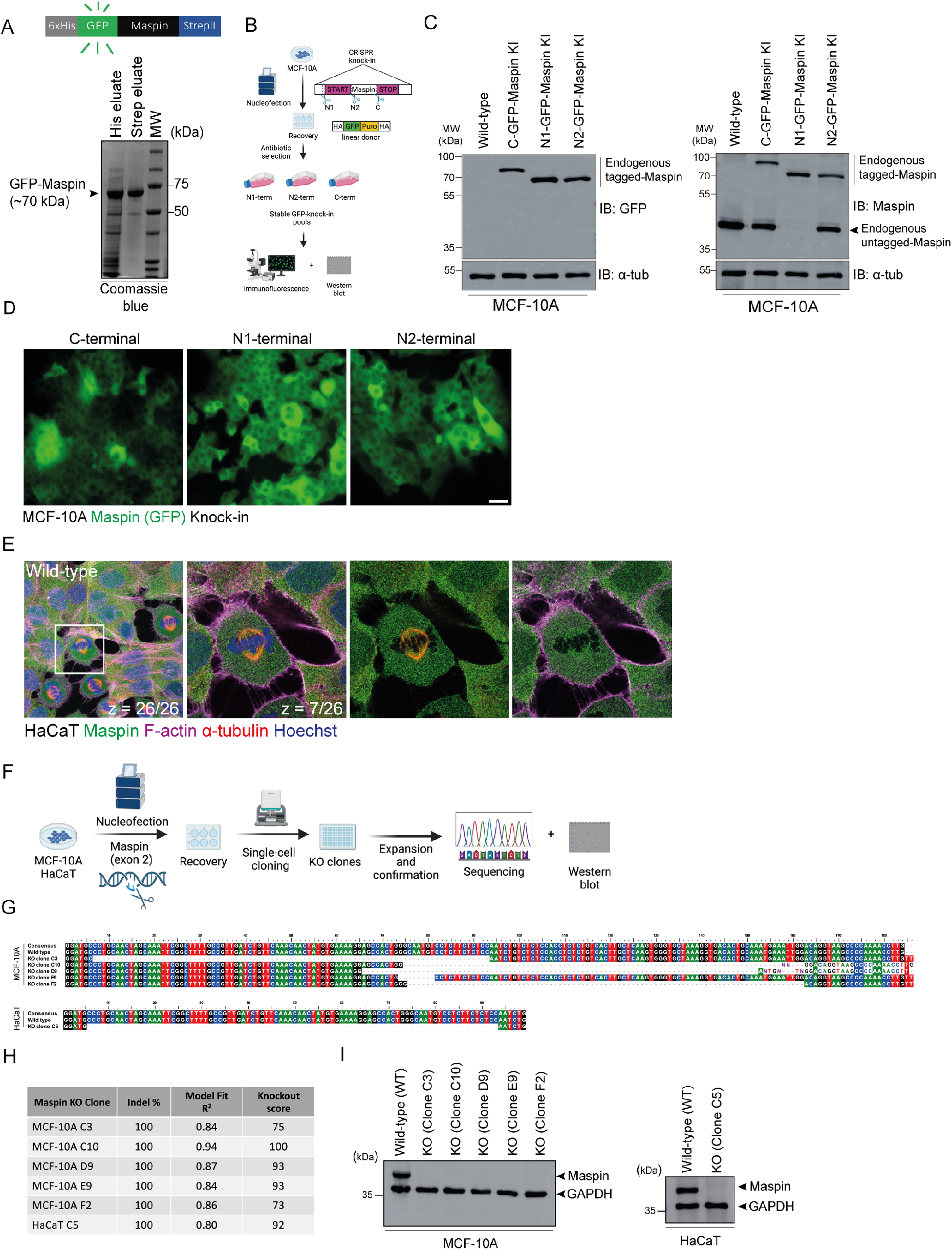
(A) Purified recombinant Maspin analyzed by SDS-PAGE followed by Coomassie blue staining; **(B)** Workflow of CRISPR/Cas9 knock-in strategy to insert a GFP tag into the Maspin locus of MCF-10A cells. Stable pools were selected with puromycin; HA = amplified homology arms, Puro = puromycin resistance sequence; **(C)** Western blot analysis for Maspin in MCF-10A knock-in pools (C-terminal, N1-terminal and N2-terminal) generated as described in B. The same cell lysates were probed using an anti-GFP antibody (left panel) or anti-Maspin antibody (right panel). Anti-α- tubulin antibody was used as a loading control; **(D)** Representative images of endogenously-tagged Maspin pools monitored by fluorescence live imaging. Scale bar: 20 µm; **(E)** Co-staining of Maspin, F- actin and α-tubulin in HaCaT cells; **(F)** Workflow of CRISPR/Cas9 knock-out strategy to target exon 2 of Maspin in MCF-10A and HaCaT cells. After 48 h of recovery from nucleofection, single cell cloning was performed and clones were further expanded. Edited regions were sequenced and protein levels assessed by Western blot; **(G)** Sequence alignment for all knock-out clones obtained in (F). Base deletions are depicted as dashed lines and base insertions as non-shaded letters; **(H)** Inference of CRISPR Edits (ICE) report for all knock-out clones employed in this study; **(I)** Western blot analysis for all KO clones using an anti-Maspin antibody. GAPDH was used as a loading control.

**Supplementary Figure 2.**
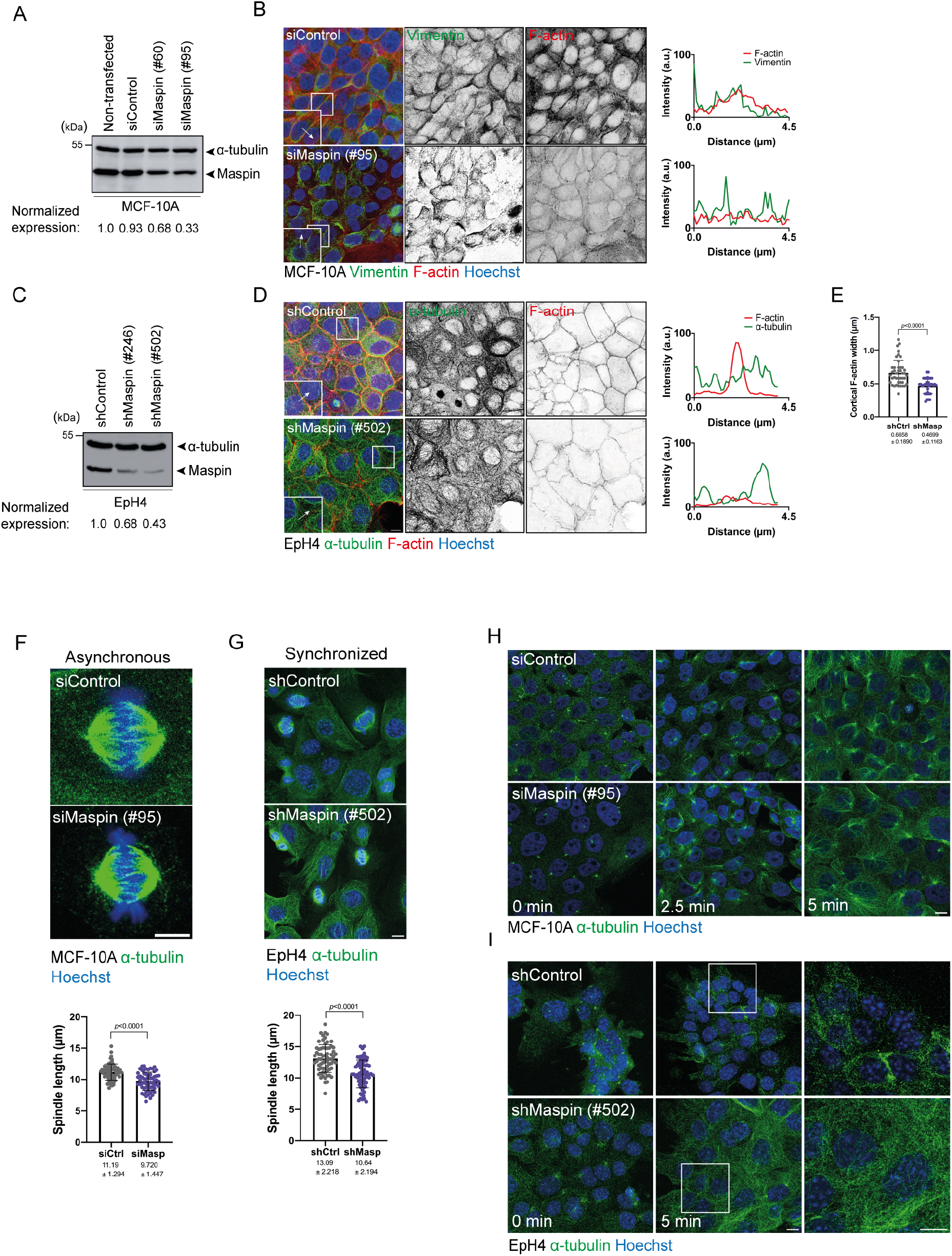
(A) Maspin protein levels of non-transfected or transiently transfected MCF- 10A cells with siControl, siMaspin (#60) and siMaspin (#95). Anti-α-tubulin antibody was used as a loading control; **(B)** Co-staining of F-actin and vimentin for MCF-10A cells transiently transfected as described in (A). Scale bar: 10 µm; **(C)** Maspin expression levels of EpH4 cells transduced with shControl, shMaspin (#246) and shMaspin (#502). Anti-α-tubulin antibody was used as a loading control; **(D)** Co-staining of F-actin and α-tubulin for EpH4 cells transduced as described in (C). Scale bar: 10 µm. Fluorescence intensity profiles refer to corresponding arrows in images; **(E)** Cortical F-actin width for EpH4 cells expressing shControl (n = 38 cells) or shMaspin (n = 40 cells); **(F)** Immunofluorescence for α-tubulin in asynchronous MCF-10A cells at metaphase. Spindle length was measured across pole-to-pole distance for siControl (n = 61 cells) and siMaspin (n = 60 cells). Scale bar: 5 µm; **(G)** Immunofluorescence for α-tubulin in thymidine-synchronized EpH4 cells at metaphase. Spindle length was measured across pole-to-pole distance for shControl (n = 68 cells) and shMaspin (n = 70 cells). Scale bar: 10 µm; **(H)** Microtubule regrowth assay for MCF-10A transfected with siControl or siMaspin (#95); **(I)** Microtubule regrowth assay for EpH4 stably expressing shControl or shMaspin (#502). Scale bar: 10 µm.

**Supplementary Figure 3.**
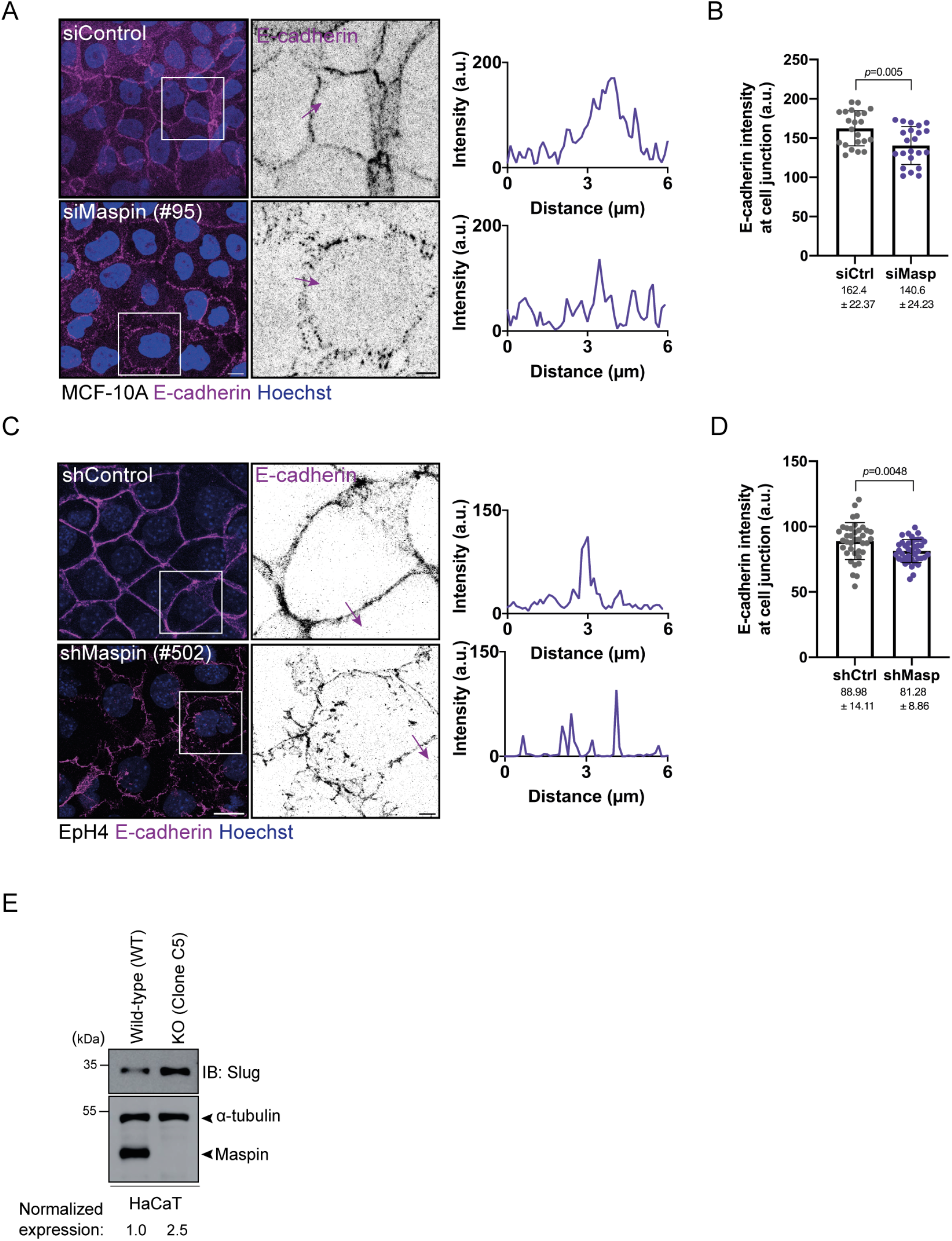
(A) Immunofluorescence analysis of E-cadherin in MCF-10A cells transiently transfected with siControl or siMaspin (#95). Scale bar: 10 µm; **(B)** E-cadherin intensity at cell junction (a.u.) in MCF-10A cells transiently transfected with siControl (n = 22 cells) and siMaspin (n = 23 cells); **(C)** Immunofluorescence analysis of E-cadherin in EpH4 cells stably expressing shControl or shMaspin (#502); **(D)** E-cadherin intensity at cell junction (a.u.) in EpH4 cells stably expressing shControl (n = 40 cells) or shMaspin (n = 38 cells); **(E)** Expression levels of Slug in HaCaT KO clone C5 *versus* WT analyzed by Western blot. Anti-α-tubulin antibody was used as a loading control.

